# A Self-Assembling Immune-Featured Osteosarcoma Patient/PDX Derived Organoid Model and Biobank for Personalized Immune Therapy

**DOI:** 10.1101/2024.08.11.607471

**Authors:** Haoran Mu, Yining Tao, Jinzeng Wang, Xin He, Qi Zhang, Weixi Chen, Bing Yao, Sen Ding, Xiyu Yang, Liyuan Zhang, Hongsheng Wang, Dongqing Zuo, Jiakang Shen, Mengxiong Sun, Haoyu Wang, Ming Jiao, Xinmeng Jin, Yinhui Jin, Youzhi Liang, Yuyan Gong, Winfred Mao, Qian Liu, Zhuoying Wang, Yu Lv, Jing Xu, Tao Zhang, Yuqin Yang, Jun Lin, Fred J. Asward, James D. Joseph, Mingxi Li, Zhengdong Cai, Wei Sun, Liu Yang, Yingqi Hua

## Abstract

Osteosarcoma (OS) exhibit intra- and inter-heterogeneity, complicating the exploration of effective therapeutic strategies. Traditional *in vitro* and *in vivo* models are limited in inheriting biological and genomic heterogeneities of OS patients, even in inheriting the features on tumor microenvironment. The prolonged generation time of current models makes the drug development of OS slow and is not suitable to clinically rapid timing. Here, we introduce methods for generating and biobanking patient/PDX-derived osteosarcoma organoids (OS PD(X)Os) that recapitulate the histological, biological and genomic features of their paired OS patients. OS PD(X)Os can be generated quickly with high reliability *in vitro* or transplanted to immunodeficient mice. We further demonstrate an immune-featured OS PD(X)O (named iOS) model and its method for testing personalized chemotherapy response, personalized immune therapeutic strategy and target drug development, such as a novel PRMT5^MTA^ inhibitor ARPN2169 on MTAP-deleted OS. Our studies show that iOS models maintain many typical features of OS and could be rapidly employed to investigate patient-specific therapeutic strategies. Additionally, our biobank establishes a rich resource for basic, translational and even clinical OS researches.

## Introduction

Osteosarcoma (OS) is considered as a primary malignant bone tumor, frequently affecting individuals in their early years of life^1–3^. Patients diagnosed with OS often confront a daunting prognosis, compounded by limited treatment options aimed at improving their survival after first-line therapy resistance. dThe median survival of OS patients is approximately 70%, with no significant improvement over the past four decades. In our previous study^4^, molecular heterogeneity among OS has been appreciated to contribute poor outcomes. Developing new models inherited OS heterogeneity for prompt testing of personalized therapy strategies for OS remain challenges in both clinical and pre-clinical stages.

Several models have contributed to understanding the biological mechanisms and drug developments of OS, but they have limitations. Traditional *in vitro* OS models could not maintain various cellular subgroups and their typical biological features in both monolayer and tumor sphere cultures. Meanwhile, due to the rare incidence of osteosarcoma ^5^, the conventional models acknowledged for their scarcity inherently lack complex genetic features. Patient-derived xenograft (PDX) models^6,7^, where dissociated tumor cells from patients are directly injected into immunodeficient mice, are believed to preserve these crucial biological features of OS more effectively^8,9^. However, PDX models do not fully inherit the immune microenvironment characteristics of osteosarcoma and exhibit a prolonged latency period for model establishment. Using traditional models or PDX models can have some utilities at the pre-clinical stages; but their development cycles are lengthy. We still need to develop models that accommodate shorter and faster experimental cycles to fit OS clinical diagnostics and treatments.

With the advancement of organoid researches, numerous models capable of simulating tissue development and organ functions have emerged, driving significant progress in physiology and oncology^10,11^. These advancements have predominantly targeted epithelial tumors, hematologic malignancies, and associated tissues^12–14^. Recent studies have demonstrated that tissue chunks in some tumor types can be cultured *in vitro* to form organoid models and biobank, ostensibly preserving original tissue characteristics^15^. Organoids have been successfully established in many tumors for large-scale high-throughput screening^16,17^. As of now, organoid development in oncology has progressed from simple drug or drug target screening in tumor cells to exploring or targeting the formation of distinct tumor immune microenvironments^18,19^. This shift aims to advance personalized clinical diagnosis and treatment. However, organoid models lacking complete immune features cannot accurately identify drugs and targets, especially immune-related therapies^18^. However, progress in constructing stromal tumor organoids with complete immune features, particularly for bone-related tumors like OS, has been slow^20,21^. Bone tumors, particularly OS, not like soft tissue sarcoma^22–24^, present challenges due to the high density and hardness of the bone matrix, making shredding or digestion extremely difficult. Additionally, irregular tumor-driven osteogenesis further complicates digestion, resulting in dense bone debris within cell suspensions. This complicates accurate cell viability measurements using high-throughput technologies and impacts the success rate of dissociating single-cell suspensions, even by using tumor chunks. These challenges not only hinder the construction of organoids for bone-related tumors but also impede research into their microenvironment and studies relying on single-cell analysis, such as single-cell sequencing.

Here, based on our previous SGH-OS clinical OS cohort^4^, we report a method to rapidly generate a self-assembling single-cell osteosarcoma patient/PDX-derived organoid (OS PD(X)O) model in a defined culture medium. Established OS PD(X)O models can complete an experimental cycle in approximately 7 to 10 days. We also generate a live biobank of OS PD(X)Os and performed histological staining, molecular and genomic analyses to identify the intra- and inter heterogeneity, which are inherited from their paired patient/PDXs. OS PD(X)Os can be xenografted in the immunodeficient mouse. Based on this biobank, we introduce a method to generate an immune-feature osteosarcoma organoid (iOS) model. We further show that iOS models can be employed to test response to standard chemotherapy of OS, as well as immunotherapy and personalized therapy exploration based on their biological features, such as PRMT5^MTA^ inhibitor APRN2169 on iOS model carried MTAP deletion and KRAS mutation, on a clinically relevant timescale. Together, these results highlight the potential functions of our iOS model and biobank for basic, translational and even clinical researches and for personalized therapies.

## Methods

### Study approval

Our study was approved by the Institutional Research Ethics Committee of Shanghai General Hospital, Shanghai Jiao Tong University School of Medicine (2021KY103). The specimens of patients or patient-derived xenografts (PDX) were derived from newly diagnosed osteosarcoma (OS) patients with written consents from Shanghai General Hospital, including clinical information. Pathological diagnoses of all the OS patients admitted in Shanghai General Hospital were independently examined by three pathologists.

### Preparation of storage solution (OS-Preservator) and washing solution (OS-Cleanser)

Each 50 ml of OS-Preservator was composed of fetal bovine serum (FBS) (2%, #10099141C, *Gibco*), penicillin-streptomycin solution (1%, #15140122, *Gibco*), HEPES (10 mmol/L, #15630080, *Gibco*) and OS-Preservator was then brought up to 50 ml with HBSS (#14170161, *Gibco*). Once the OS-Preservator was prepared, it was aliquoted using 15 mL sterile centrifuge tubes (#430053, *Corning*), 5 mL per tube, and stored at 4°C. OS-Preservator was valid for up to 1 month. Each 50 ml of OS-Cleanser was composed of penicillin-streptomycin solution (1%) and OS-Cleanser was then brought up to 50 ml with PBS (#21-040-CVR, *Gibco*). OS-Cleanser was prepared for immediate use.

### Preparation of dissociation solution (OS-Dissolver) and dissociation termination solution (OS-Terminator)

Each 5 ml of OS-Dissolver was composed of collagenase II (125 CDU/mL, #V900892, *Sigma-Aldrich*), DNase I (40 Kunitz units/mL, #DN25, *Sigma-Aldrich*), penicillin-streptomycin solution (0.1%), FBS (5%) and OS-Dissolver was then brought up to 5 ml with RPMI 1640 medium (#SH30809.01B, *Hyclone*). OS-Dissolver was prepared for immediate use. Define the unit of collagenase (collagenase II) activity, CDU, by protease activity: at 37°C and pH 7.5, 1 CDU of protease releases 1 μmol of L-leucine from collagen in 5 hours. Define the unit of DNase I activity, 1 Kunitz unit, as the amount of enzyme that causes an increase in absorbance of 0.001 A260nm/min/mL due to the degradation of highly polymerized DNA at 25°C (0.1 M NaOAc, pH 5.0). Each 50 ml of OS-Terminator was composed of penicillin-streptomycin solution (1%), FBS (10%) and OS-Terminator was then brought up to 50 ml with Dulbecco’s Modified Eagle Medium F12 (DMEM/F12) (#11320033, *ThermoFisher*). OS-Terminator was stored at 4°C for up to 1 month. OS-Dissolver and OS-Terminator were prepared in *SeekGene* (*SeekGene Corporation*, China).

### Preparation of culture medium (OS-Nurturer) and cryopreservation solution (OS-CytoPreserver)

Each 50 ml of OS-Nurturer was composed of penicillin-streptomycin solution (1%), FBS (10%), GlutaMAX^TM^ supplement (1%, #35050061, *Gibco*), N-2 supplement (1.2%, #17502048, *ThermoFisher*), MEM Non-Essential Amino Acids (NEAA) (1.2%, #11140050, *ThermoFisher*), B-27^TM^ supplement (1.2%, #17504044, *ThermoFisher*), Insulin, Transferrin, Selenium Solution (ITS-G) (1.2%, #41400045, *Gibco*), Ascorbic acid (100 μmol/L, #200-066-2, *Merck*), ROCK inhibitor Y-27632 (10 μmol/L, #10035-04-8, Merck), human recombinant EGF protein (20 ng/mL, #AF-100-15, *PeproTech*), human recombinant bFGF protein (40 ng/mL, #HY-P7331A, *MedChemExpress*), human recombinant IGF-1 protein (20 ng/mL, # HY-P70783, *MedChemExpress*), human recombinant TGF-β3 protein (10 ng/mL, #HY-P7120, *MedChemExpress*), Adenine (188 μmol/L, #HY-B0152, *MedChemExpress*), ALK inhibitor A 83-01 (0.5 μmol/L, #HY-10432, *MedChemExpress*), selective p38 MAPK inhibitor SB 202190 (10 μmol/L, #HY-10295, *MedChemExpress*), Dexamethasone (10^-5^ mol/L, #HY-14648, *MedChemExpress*), human recombinant Wnt3a protein (500 ng/ml, #HY-P70453A, *MedChemExpress*) and OS-Nurturer was then brought up to 50 ml with DMEM. OS-Nurturer was sterilized by filtration using a 0.22 μm syringe filter (#SLGP033RS, *Millipore*). OS-Nurturer was stored at 4°C for up to 3 weeks. Each 10 ml of OS-CytoPreserver was composed of Y-27632 (10 μmol/L) and OS-CytoPreserver then brought up to 10 ml with serum-free cell cryopreservation solution (#C40100, *NCM Biotech*). OS-CytoPreserver was prepared for immediate use.

### Single-cell OS patient/PDX derived organoid (PD(X)O) suspension preparation from osteosarcoma patients or PDXs

Under sterile conditions, retrieve tumor tissue from osteosarcoma patients or PDXs. Store it at 4°C using OS-Preservator and promptly transport it to the laboratory, ideally within 2 hours. Assess tissue firmness to determine the necessity of dilution with OS-Dissolver. Utilize OS-Cleanser to meticulously cleanse the tissue, eliminating non-tumor tissue, necrotic tissue, blood clots, and other contaminants. Cut the tissue into small fragments within clean OS-Cleanser, ensuring each piece measures approximately 1.5-3 mm³ to prevent excessive bone shard production that may impact experimental integrity. Employ OS-Dissolver at 37°C for approximately 15 minutes to dissociate the tissue. Monitor the dissociation process under a microscope until numerous single cells are visible. Mix OS-Dissolver with at least three times its volume of OS-Terminator, gently pipetting to ensure thorough blending. Filter the resultant cell suspension through a sterile 40 μm cell strainer (#431750, *Corning*) to eliminate residual tissue debris. Centrifuge (1500 G) to discard the supernatant. Carefully resuspend the cell pellet in PBS, then centrifuge (1500 G) again to remove any remaining OS-Terminator.

### Self-assembly and cultivation of OS PD(X)O

Count the harvested single-cell OS PD(X)O suspension. At room temperature (∼25°C), adjust the concentration of the single-cell OS PD(X)O suspension to 2 × 10^5^ cells/50 μl using OS-Nurturer, and distribute 50 μl into each well of a 96-well clear round-bottom ultra-low attachment microplate (#7007, *Corning*). Centrifuge the 96-well plate at 500 G for 3 minutes at room temperature, carefully remove it, and gently place it on the workstation. Prepare a self-assembling matrix by combining Matrigel^®^ (#356231, *Corning*) with OS-Nurturer in a 3:2 ratio at 4°C or on ice, and gently add 75 μl per well over the centrifuged single-cell suspension at 4°C or on ice. After addition, gently transfer the 96-well plate to a cell culture incubator (37°C, 5% CO_2_) and allow it to sit undisturbed for 30-60 minutes to facilitate thorough mixing and cross-linking of the self-assembling matrix with OS PD(X)O. Once the cross-linking is complete, remove the plate and gently add 100-150 μl of OS-Nurturer per well at room temperature. Return the plate to the cell culture incubator (37°C, 5% CO_2_) and replace the OS-Nurturer every 1-2 days. Monitor the growth of OS PD(X)O using bright-field microscopy until organoid-like structures with a diameter of 1000-1500 μm are observed.

### Passaging, cryopreservation and resuscitation of OS PD(X)O organoid

Using a 4°C sterile PBS solution (200 μl per well), gently aspirate and collect the OS PD(X)O organoid models designated for passaging to dissolve the Matrigel^®^. Transfer the dissolved structures to a 15 ml centrifuge tube and let them stand at 4°C for 15 minutes. Add 1-2 ml of sterile PBS solution and gently mix by pipetting the organoid-like structures. Centrifuge at 1500 G for 5 minutes at room temperature, then carefully remove the supernatant. Add 1-2 ml of OS-Dissolver, gently pipette to ensure thorough mixing, and incubate at cell culture incubator (37°C, 5% CO_2_) for 10 minutes or until the organoid-like structures dissociate into single cells. Stop the digestion reaction by adding OS-Terminator at three times the volume of OS-Dissolver used. Collect the single-cell suspension by centrifugation at 1500 G for 5 minutes at room temperature. Resuspend the cell pellet in OS-Nurturer, and proceed with cultivation after passaging OS PD(X)O through 2-3 generations for expansion. For cryopreservation, using a 4°C sterile PBS solution (200 μl per well), gently aspirate and collect the OS PD(X)O organoid intended for passaging to dissolve the Matrigel^®^. Transfer to a 15 ml centrifuge tube and let stand at 4°C for 15 minutes. Add 1-2 ml of sterile PBS solution and gently mix by pipetting the organoid-like structures. Centrifuge at 1500 G for 5 minutes at room temperature. Resuspend the organoid cells in OS-CytoPreserver at 0.5-1 ml per well, and gradually cool down to 4°C (∼1-2 hours) before transferring to liquid nitrogen for long-term storage. For revival from liquid nitrogen, prepare sterile 37°C water in advance. Quickly thaw the OS PD(X)O from the cryovial in a 37°C water bath. Centrifuge at 1500 G at room temperature for 5 minutes, carefully remove the supernatant, and resuspend the OS PD(X)O organoid single-cell suspension in OS-Nurturer. Observe cell morphology under a microscope, count the cells, and proceed with cultivation.

### Co-cultivation of OS PD(X)O with peripheral blood mononuclear cells (PBMC)

Specialized co-cultivation medium was prepared based on OS-Nurturer, supplemented with human recombinant IL-4 (5 ng/mL, #200-04, *PeproTech*) and human recombinant GM-CSF (5 ng/mL, #300-03, *PeproTech*). PBMCs were purchased from *Milestone^®^* (#PB025C, *TPCS*, *Milestone^®^ Biotechnologies*), the donors were obtained informed consent. If PBMC pre-activation is required, cells were treated with FITC anti-human CD3 antibody (10 μg/mL, #300306, *BioLegend*) in a 96-well plate. Following this, CD28 Monoclonal Antibody (5 μg/ml, #16-0289-85, *Invitrogen*) and Human Recombinant IL-2 (1 μl/10 ml, #200-02, *PeproTech*) were incubated with the cells at 37°C for 2 hours. Mix the dissociated single-cell OS PD(X)O organoid suspension with treated PBMCs in a specified ratio, and following the culture method for self-assembling OS PD(X)O organoid models, perform self-assembling matrix and cultivation, but the culture medium used is RPMI 1640 mixed with 2× Co-cultivation medium in a specified ratio. Change the medium every two days, and if anti-human PD-1 antibody (#A2002, *Shlleck*) or PRMT5^MTA^ inhibitor ARPN2169 (this study, *APEIRON*) is added to the culture conditions, also change the medium every two days. The cultivation period is 7-10 days.

### 3D cell viability assay

Cell viability was measured by *CellTiter-Lumi™ luminescent cell viability assay kit* (#C0065S, *Beyotime*). The preparation and protocol of cell viability assay kit were according to *Beyotime* official website and abbreviated as follows. Allow the cell culture plate to equilibrate at room temperature for 10 minutes (not exceeding 30 minutes). Add 100 μl of CellTiter-Lumi™ luminescent assay reagent per well for a 96-well plate. Gently shake at room temperature for 2 minutes to facilitate cell lysis. Incubate at room temperature for 10 minutes to stabilize the luminescent signal. Perform luminescent detection was used by SpectraMax M3 Microplate Reader (*Molecular Devices*). Calculate relative cell viability directly from the luminescent readings or determine ATP content using an ATP standard curve to assess relative cell viability based on ATP levels. Nuclease free ATP was purchased from *Beyotime* (#D7378, *Beyotime*).

### Flow cytometry

Centrifuge the single-cell suspension and discard the supernatant. Resuspend the cells in 300-500 μl of FACS staining buffer (#00-4222-26, *Invitrogen*), add 1 μl of Blocking buffer (#B001T06F01, *Invitrogen*), gently vortex to mix, and surface stain at 4°C for 15 minutes with 1 μl of antibody (1:500 dilution, various antibodies can be mixed). Vortex thoroughly to ensure even staining, then wash twice with 1 ml of FACS staining buffer, vortexing between washes. Centrifuge, discard the supernatant, and fix with 300 μl of PFA (final concentration at 2-4%) at 4°C for 30 minutes. Wash with PBS, store in the dark at 4°C, and analyze within 3 days. The antibodies used include PerCP anti-human CD45 antibody (#304025, *BioLegend*), FITC anti-human CD3 antibody (#317305, *BioLegend*), APC anti-human CD4 antibody (#317415, *BioLegend*), PE anti-human CD8 antibody (#344705, *BioLegend*), and BV605 anti-human PD-1 antibody (#367425, *BioLegend*). For flow cytometry compensation adjustments, resuspend splenic lymphocytes in 300-500 μl of FACS staining buffer. Add 1 μl of the relevant antibody (1:500 dilution), mix gently by vortexing, and incubate at 4°C for 30 minutes. Proceed with washing twice with 1 ml of FACS staining buffer, centrifuge, discard the supernatant, fix with 300 μl of PFA (final concentration at 2-4%) at 4°C for 30 minutes, wash with PBS, store in the dark at 4°C, and analyze by LSR Fortessa flow cytometry (*BD Biosciences*, USA) within 3 days.

### Mouse xenografts of osteosarcoma

6-8-week-old female C3H mice (Strain: C3H/HeNCrl; Charlers river, USA/China) and 6-8-week-old female Nude mice (Strain: BALB/cJGpt-foxn1nu/Gpt; Charlers river, USA/China) were used in this study. DuNN MTAP-del and its culture were according to our previous study (https://dx.doi.org/10.2139/ssrn.4881940). All mice were maintained under SPF conditions in a controlled environment of 20-22℃, with a 12/12h light/dark cycle, 50-70% humidity. All animal experiments were performed by following the protocols in accordance with the guidelines of Laboratory Animal Center of Shanghai General Hospital. The Clinical Center Laboratory Animal Welfare & Ethics Committee of Shanghai General Hospital, Shanghai Jiao Tong University School of Medicine, approved all animal protocols used in this study. The OS mouse xenograft was developed by intramedullary injection with OS cells (5×10^5^ cells in 25ul PBS supplemented with 10% FBS) into the marrow space of the proximal tibial of C3H or Nude mice with a 27-gauge needle, as our previous studies^25,26^. OS progression in the mice was monitored by measuring the tumor volume, calculated by the following equation: Volume = Length×Width^2^/2. The tumor burden was monitored by following the tumor volume and the maximal tumor size is 2 cm, which was permitted by the Clinical Center Laboratory Animal Welfare & Ethics Committee of Shanghai General Hospital, Shanghai Jiao Tong University School of Medicine. The maximal tumor size was confirmed to be not exceeded in our study and the survival curves were produced according to the survival of mice and the time to reach maximum tumor size.

### Establishment and passaging of PDX models

Establishment and passaging of PDX models were as our previous study ^4^. Approximately 100 mg of tissue was placed in a 15 ml polypropylene tube containing serum-free DMEM and transported to the laboratory on wet ice. After washing and sectioning, tumor fragments measuring 3–5 mm were implanted into the flanks of NSG mice. Tumor size and body weight were monitored twice weekly, and tumor volume was calculated using the formula: Length × Width^2^ × 0.5 mm^3^. All experimental procedures involving animal subjects were conducted in accordance with the ethical guidelines and approved by the Shanghai General Hospital Animal Care and Use Committee.

### Immunohistochemical (IHC) and Hematoxylin & Eosin (H&E) staining

IHC staining was performed on representative sections from formalin-fixed and paraffin-embedded tissue using the mentioned antibodies at the indicated concentrations below. The antibodies were including MKI67 (1:500, #GB151499-100, *Servicebio*), SATB2 (1:500, #GB111449-100, *Servicebio*) and VIM (1:500, #GB121308-100, *Servicebio*). H&E staining was performed as our previous study ^27^. IHC and H&E staining were performed in *Servicebio Corporation* (*Servicebio*, China).

### Whole genome sequence (WGS) and analysis

The genomic DNA was extracted based on the CTAB method. About 200 mg tissue after ground to powder by liquid nitrogen was transferred to a preheated (65℃) 2.0 ml tube with appropriate amount of CTAB lysis buffer (#MC1411, *Promega*) and was mixed by vortex. The tube was incubated at 65℃ for 60 minutes, then was centrifuged at 10,000 rpm at room temperature (RT) for 5 minutes. The supernatant was extracted with one volume of phenol/chloroform/isopentanol (25:24:1) followed by centrifuging at 10,000 rpm at RT for 10 minutes in a new tube. 2/3 volume of precooled(-20℃) isopropanol was added into the tube and put at -20℃ for more than 2 hours to precipitate the DNAs, followed by centrifuging at 12,000 rpm for 15 minutes at RT. 75% ethanol was added to wash the pellet and removed by centrifuging, then the DNA pellet was air-dried for 3-5 minutes. The pellet was dissolved by 30-200 μl TE buffer (#12090015, *ThermoFisher*) for further study. After DNA extraction, 1 μg genomic DNA was randomly fragmented by Covaris Focused-ultrasonicators (*Covaris*), followed by fragments selection by Agencourt AMPure XP-Medium kit to an average size of 200-400 bp. Selected fragments were end repaired and 3’adenylated, then the adaptors were ligated to the ends of these 3’adenylated fragments. The products were amplified by PCR and purified by the Agencourt AMPure XP-Medium kit (#A63882, *Beckman*). The purified double stranded PCR products were heat denatured to single strand, and then circularized by the splint oligo sequence. The single strand circle DNA (ssCir DNA) were formatted as the final library and qualified by quality control (QC). The final qualified libraries were sequenced by BGISEQ-500 (*BGI Genomics*). ssCir DNA molecule formed a DNA nanoball (DNB) containing more than 300 copies through rolling-cycle replication. The DNBs were loaded into the patterned nanoarray by using high density DNA nanochip technology. Finally, pair-end 100 bp reads were obtained by combinatorial Probe-Anchor Synthesis (cPAS). WGS was prepared and performed at *Benagen* (*Wuhan Benagen Technology Co., Ltd.*). Analysis methods are as follows: Xgenome was used to remove potential contaminated mouse reads from WGS data after quality control. The filtered reads were aligned to the UCSC hg19 reference sequence with Burrows-Wheeler Aligner (bwa mem). PCR duplicates were removed by Picard, and the BAM files were then indexed by Samtools. Base quality score recalibration was performed by the BaseRecalibrator and ApplyBQSR tools from the Genome Analysis Toolkit (GATK) according to GATK best practices. Somatic variants including single nucleotide variants (SNVs) and small indels were detected using Mutect2 in GATK on processed genome data with tumor-only model. Annotation of variants was carried out by Annovar on the Refseq gene model. Variants in the non-coding regions (upstream, downstream, intergenic, intronic, ncRNA, UTR5, UTR3, etc.) were excluded from the analyses. Germline variants were filtered by using the 1000 Genomes, Exome Aggregation Consortium, NHLBL Exome Sequencing Project (ESP6500) and Genome Aggregation Database (gnomAD). Mutation landscape was generated by using maftools. The CNVkit was applied to calculate somatic copy number alterations (SCNAs) with default parameters based on WGS data. Clustering of SCNAs by chromosomes was performed in R based on Euclidean distance using Ward’s complete method by the pheatmap package.

### Bulk RNA sequencing (RNA-seq) and analysis

RNA purity of specimens was checked using the kaiaoK5500 Spectrophotometer (*Kaiao*). RNA integrity and concentration was assessed using the RNA Nano 6000 Assay Kit of the Bioanalyzer 2100 system (#5067-1511, *Agilent Technologies*). A total amount of 2ug RNA per sample was used as input material for the RNA sample preparations. Sequencing libraries were generated using NEBNext Ultra RNA Library Prep Kit for Illumina (#7770, *New England Biolabs*) according to the manufacturer’s protocol and index codes were added to attribute sequences to each sample. The clustering of the index-coded samples was performed on a cBot cluster generation system using HiSeq PE Cluster Kit v4-cBot-HS (#PE-401-4001, *Illumina*) according to the manufacturer’s instructions. After cluster generation, the libraries were sequenced on DNBSEQ-T7 platform (*BGI Genomics*) in *Benagen* (*Wuhan Benagen Technology Co., Ltd.*) and 150 bp paired-end reads were generated. The raw data were first processed with FastQC to filter out adapters and low-quality sequences. Pair-end reads were aligned to human GRCh38 genome using STAR. Reads with good mapping quality (MAPQ > 30) that aligned to genomic exons were counted using featureCounts (GRCh38) to generate a table with counts for each gene. Principal Component Analysis (PCA) was performed using the R package stats. Differential gene expression analysis was performed using the R package DESeq2 using the IfcShrink function. Genes with fold-change ≥ 2.00, probability ≥ 0.80 and false discovery rate P value (FDR) < 0.05 were considered significantly differentially expressed. Enrichment analyses for differentially expressed genes were performed using the R package clusterProfiler. Gene set enrichment analysis (GSEA, http://www.gsea-msigdb.org/gsea/) was performed on list of genes ranked from high to low DESeq2 estimated fold-change using the GSEAPreRanked function with enrichment statistic classic, 1000 permutations and normalized P value < 0.05. Single sample GSEA (ssGSEA) was performed by the R package GSVA using the gene sets curated. Related gene sets were downloaded from Molecular Signatures Database (MSigDB, https://www.gsea-msigdb.org/gsea/msigdb/).

### KRAS^G12C^ mutation gene probe assay and analysis

The formalinfixed, paraffin-embedded (FFPE) specimens were obtained from the osteosarcoma patient/PDXs and OS PD(X)O models. DNA extraction was performed using a TIANamp® FFPE DNA Kit (#4992524, *TIANGEN Biotech*) according to the kit’s recommended protocol. The extracted DNA was quantified using a PikoGreen dsDNA quantitative kit (#AN54L018, *Life iLab Biotech*). The pre-amplification procedure was carried out in a 25 μl reaction volume using 33.3 ng of genomic DNA, 2 × PCR Precision™ MasterMix (#G124, *Abm*), and 200 nM of the forward (5’-GTGACATGTTCTAATATAGTC-3’) and reverse primers (5’-GGATCATATTCGTCCACAAA-3’). Samples were placed in a *Mastercycler^®^* RealPlex instrument (*Eppendorf*) set at a temperature profile of 94 ℃ for 3 minutes, followed by 30 cycles of amplification (94 ℃ for 10 s, 55 ℃ for 30 s and 72 ℃ for 20 s) and a final 72 ℃ extension for 1 minute. For the limit of detection (LOD) assays, 1%, 0.1% and 0.01% VAFs samples prepared by mixing WT amplicons and SNV amplicons (WT plus SNV at a final concentration of 10 nM) were pre-amplified as above. Two microliters of the pre-amplification products were used as input to perform A-Star. The A-Star enrichment system for the target (KRAS^G12C^) consisted of 100 nM *Pf*Ago, gDNAs (forward gDNA 5’-TTTGGAGCTGATGGCG-3’,reverse gDNA 5’-TCTACGCCAGCAGCTC-3’) at a concentration 20-fold higher than that of *Pf*Ago, 200 nM of the forward and reverse primers, and 100 μM Mn^2+^, which were mixed in 2× PCR Taq MasterMix (#G888, *Abm*). Enrichment proceeded in a *Mastercycler^®^* RealPlex instrument with a temperature profile of 94 ℃ for 3 minutes, followed by 25 cycles of amplification (94 ℃ for 30 s, 55 ℃ for 30 s and 72 ℃ for 20 s) and a final 72 ℃ extension for 1 minute. The primers and probes for the target (KRAS^G12C^) were designed using *Beacon Designer’s* standard assay design pipeline and ordered as individual primers and probes from *Sangon* (*Sangon Biotech*). To validate the enriched mutant products, assays were performed using *AceQ^®^* qPCR Probe Master Mix (#Q112-02, *Vazyme Biotech*), and the results were quantified in a *StepOnePlus™* Real-Time PCR System (*ThermoFisher*) with the primer (forward primer 5’-AGGCCTGCTGAAAATGACTG-3’, reverse primer 5’-GCTGTATCGTCAAGGCACTCT-3’) and probe (mutant probe 5’-TTGGAGCTTGTGGCGTA-3’, wild-type probe 5’-TTGGAGCTGGTGGCGTA-3’, and the probe fluorophores used for mutant and wild-type DNA were FAM and VIC, respectively) sets at final concentrations of 250 nM primers and 200 nM probes. The temperature profile for amplification consisted of an activation step at 95 ℃ for 8 minutes, followed by 40 cycles of amplification (95 ℃ for 15 s and 61.5 ℃ for 40 s). To quantify the enriched mutant product, we prepared a series of standard solutions containing WT and SNV amplicon with concentrations ranging from 100 pM to 100 aM, which were quantified using the PikoGreen dsDNA quantitative kits. Then the standard curves were generated by ploting the threshold cycle (Ct) values against the logarithm of the copy numbers. The enriched sample was analyzed by *TaqMan^®^* qRT-PCR (#4673409001, *Roche*) and then the copy number of WT or SNV is calculated using its Ct value based on the standard curve. The enrichment percentage and fraction of mutant alleles mentioned in study is calculated as the ratio of the copy number of SNV/(WT + SNV).

### Statistical analysis and data processing

Details of statistical analyses of the various experiments are described in the relevant methods section. If not specified, statistical analysis was carried out using GraphPad Prism 8 software (*GraphPad Software*). After confirming that values followed a normal distribution, two-tailed Student’s t test was applied to determine the significance of differences between two groups of independent samples. Spearman’s correlation analysis was performed to determine the correlation between two group of variables. A p value < 0.05 was considered statistically significant. Details of the data points shown were described in the respective figure legends. Flow cytometry results were analyzed by FlowJo v10 (*Tree Star*). All schematic diagrams were created using *BioRender* (*BioRender.com*).

## Results

### Self-assembly of osteosarcoma patient/PDX derived organoid models

Our study established single-cell self-assembly osteosarcoma patient/PDX derived organoid (OS PD(X)O) models using specimens from OS patients or patient-derived xenograft (PDX) models. At this stage, OS PD(X)O organoid models does not manifest complete lymphocyte immune activity. While OS PD(X)O models may retain residual lymphocytes initially, these cell populations typically decline with subsequent passages. The establishment of OS PD(X)O models involves two steps: the first step is preparing single-cell suspensions, and the second step is conducting self-assembly culture (**Figure 1**). In our study, we selected fresh, highly viable human osteosarcoma tumor tissue or PDXs, removed necrotic tissue, and washed it. Using the formulation and dissociation method proposed in this study, we obtained active single-cell suspensions (**Supplementary Figure 1a**). Subsequently, the single-cell suspensions were used to construct OS PD(X)O models (**Supplementary Figure 1b**).

**Figure 1.**
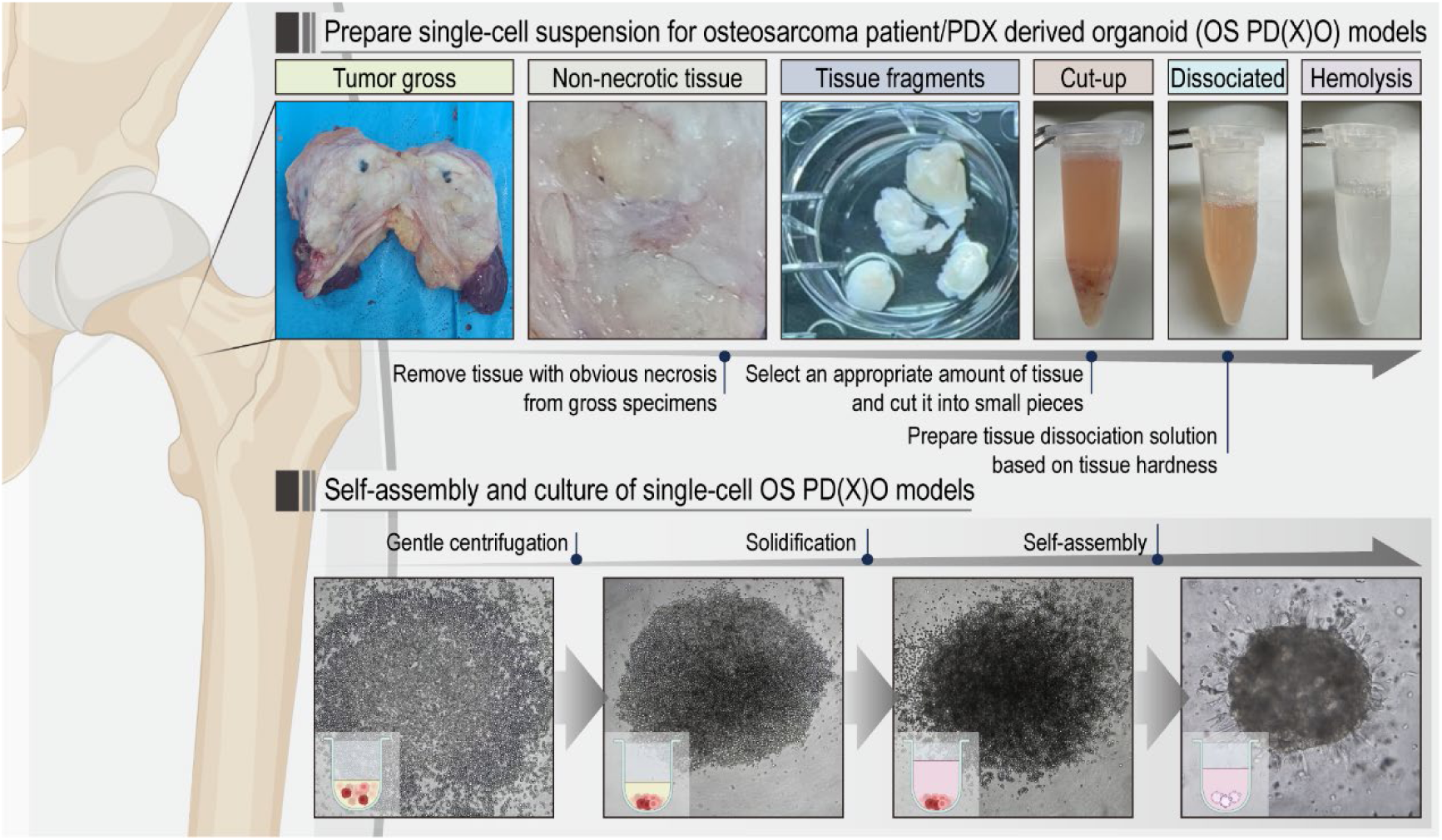
Generation of self-assembling osteosarcoma patient/PDX derived organoid (OS PD(X)O) models. See also **Supplementary Figure 1.**

We tested whether the OS PD(X)O models constructed in this study could also form subcutaneous models in animals. We further proceeded with subcutaneous tumorigenesis using corresponding PDO/PDXO constructed from Patient/PDX (**Supplementary Figure 2a**). We conducted tumorigenicity experiments using a pair of PDX/PDXO, PDXO_111. The PDXO xenograft exhibited similar growth rates to its corresponding PDX, showing no significant differences (**Supplementary Figure 2b and 2c**). The PDXO xenograft also recapitulated the molecular biology characteristics of the corresponding PDX (**Supplementary Figure 2d**). Taken together, these results demonstrate the successful establishment of our OS PD(X)O models.

### Establishment of an osteosarcoma patient/PDX derived organoid biobank

Building upon the self-assembly OS PD(X)O models, our study further established an steosarcoma patient/PDX derived organoid (OS PD(X)O) biobank from 40 osteosarcoma patients or PDX models, and this biobank contained 47 models, including 19 OS PDO models and 28 OS PDXO models (**Figure 2a**). Clinical information of paired-PD(X)O patients included gender, age, tissue lesion, primary tumor lesion and pathological subtype (**Supplementary Table 1**). In our biobank, OS PD(X)O models grew increasingly opaque spheroids with volume gradually increasing under bright-field condition (**Figure 1b, 1d and Supplementary Figure 3a**). Our OS PD(X)O models typically allowed passaging within 7 to 10 days, and due to individual differences, the growth of OS PD(X)O models exhibited varying growth rates (**Figure 1c and 1e**). In theory, each well should contain only one OS PD(X)O model. However, in our study, we found that occasionally even a small number of dispersed single cells around can grow into OS PD(X)O models. This phenomenon did not affect the functionality of our OS PD(X)O model, but it is intriguing. We initially assessed the pathological characteristics of models in the OS PD(X)O biobank. The OS PD(X)O models in our biobank inherited the pathological characteristics of the paired-patient/PDX specimens, such as large, deeply stained, and highly heterogeneous nuclei and areas resembling tumor osteogenesis (**Figure 1f and Supplementary Figure 3a**).

**Figure 2.**
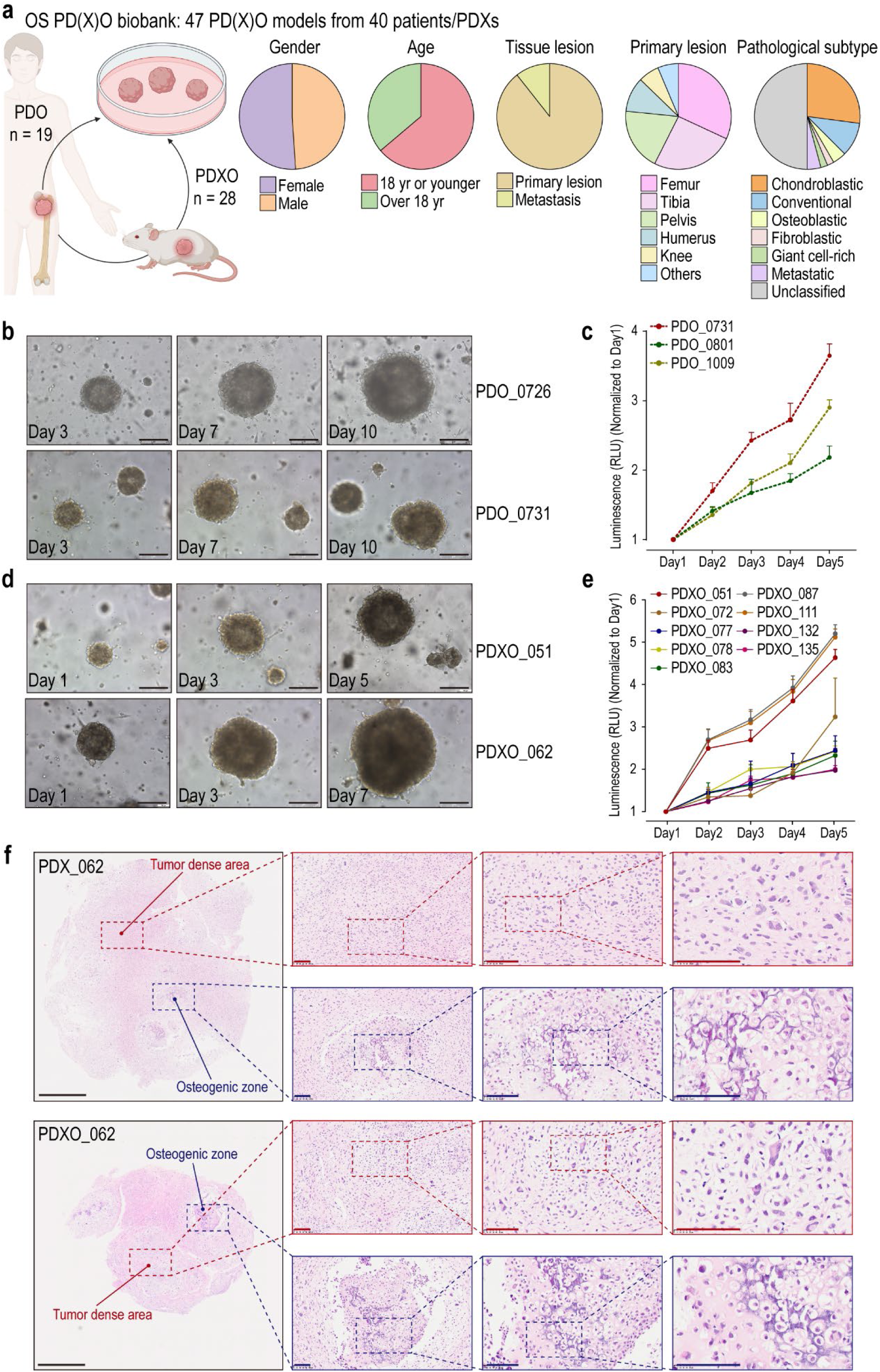
Generation of OS PD(X)O biobank that retain histological features of paired patients/PDXs. **(a)** A schematic of OS PD(X)O biobank clinical information, including gender, age, tissue lesion, primary tumor lesion and pathological subtype. **(b)** Sample brightfield images of OS PDOs. Age of OS PDOs in days is listed. Scale bars, 250 μm. **(c)** 3D cell viability assay. Quantifications of OS PDOs viability in 3 different models, corresponding for different culture days. Values represent mean ± SD (n = 4). **(d)** Sample brightfield images of OS PDXOs. Age of OS PDXOs in days is listed. Scale bars, 250 μm. **(e)** 3D cell viability assay. Quantifications of OS PDOs viability in 9 different models, corresponding for different culture days. Values represent mean ± SD (n = 4). **(f)** Sample H&E staining images of OS PDXO and paired PDX. Inheriting of intra-heterogeneity such as tumor dense area (red) and osteogenic zone (blue) Black scale bars, 250 μm. Red and blue scale bars, 100 μm. See also **Supplementary Figure 2, Supplementary Figure 3** and **Supplementary Table 1**.

We assessed whether our OS PD(X)O biobank inherited the genetic characteristics of paired patient/PDX by evaluating the transcriptomic features. We analyzed the transcriptomic profiles and employed Principal Component Analysis (PCA) to reduce the dimensionality of the paired OS PD(X)O model and patient/PDX specimen transcriptomes. The transcriptomic data distribution of each paired sample, Passage1 (P1) of OS PD(X)O and paired patient/PDX model, was similar, indicating no differences between paired samples and confirming their origin from the same population (**Figure 3a**). Based on both differential gene expression analysis (DEG) and Gene Set Variation Analysis (GSVA), we found similarities in functional enrichment of gene expression and related biological signaling pathways between each paired sample (**Figure 3b**). Further validation of molecular features was conducted using IHC, demonstrating consistency in proliferation markers MKI67 and tissue morphology, such as SATB2 and VIM, across paired samples within the biobank (**Figure 3c**). These findings suggest that OS PD(X)O models adequately preserve the molecular characteristics of their matched tissues. Further investigation was conducted to determine whether there are changes in the molecular characteristics of OS PD(X)O models during passaging. Morphologically, OS PD(X)O models showed no changes under bright-field microscopy during passaging (**Figure 3d, Supplementary Figure 4a, 4b and 4c**). PCA of transcriptomic data from passaged samples indicated similar distributions among paired samples in each passaging batch, showing no significant differences, and suggesting they originate from the same population (**Figure 3e**). Correlation analysis of the transcriptomes indicated similarity among paired samples across passaging batches, with minimal differences observed compared to their matched tissues (**Figure 3f and Supplementary Figure 4d**). Additionally, based on differential gene expression (DEG) and Gene Set Variation Analysis (GSVA), it was found that passaged samples exhibit similarity in functional enrichment of gene expression and related biological signaling pathways (**Figure 3g**). These findings suggest that cases in the OS PD(X)O biobank successfully inherit the molecular characteristics of paired tissues from patients/PDXs and maintain them throughout the passaging process. And each OS PD(X)O model also exhibits unique transcriptomic molecular features, reflecting the molecular diversity in our biobank.

**Figure 3.**
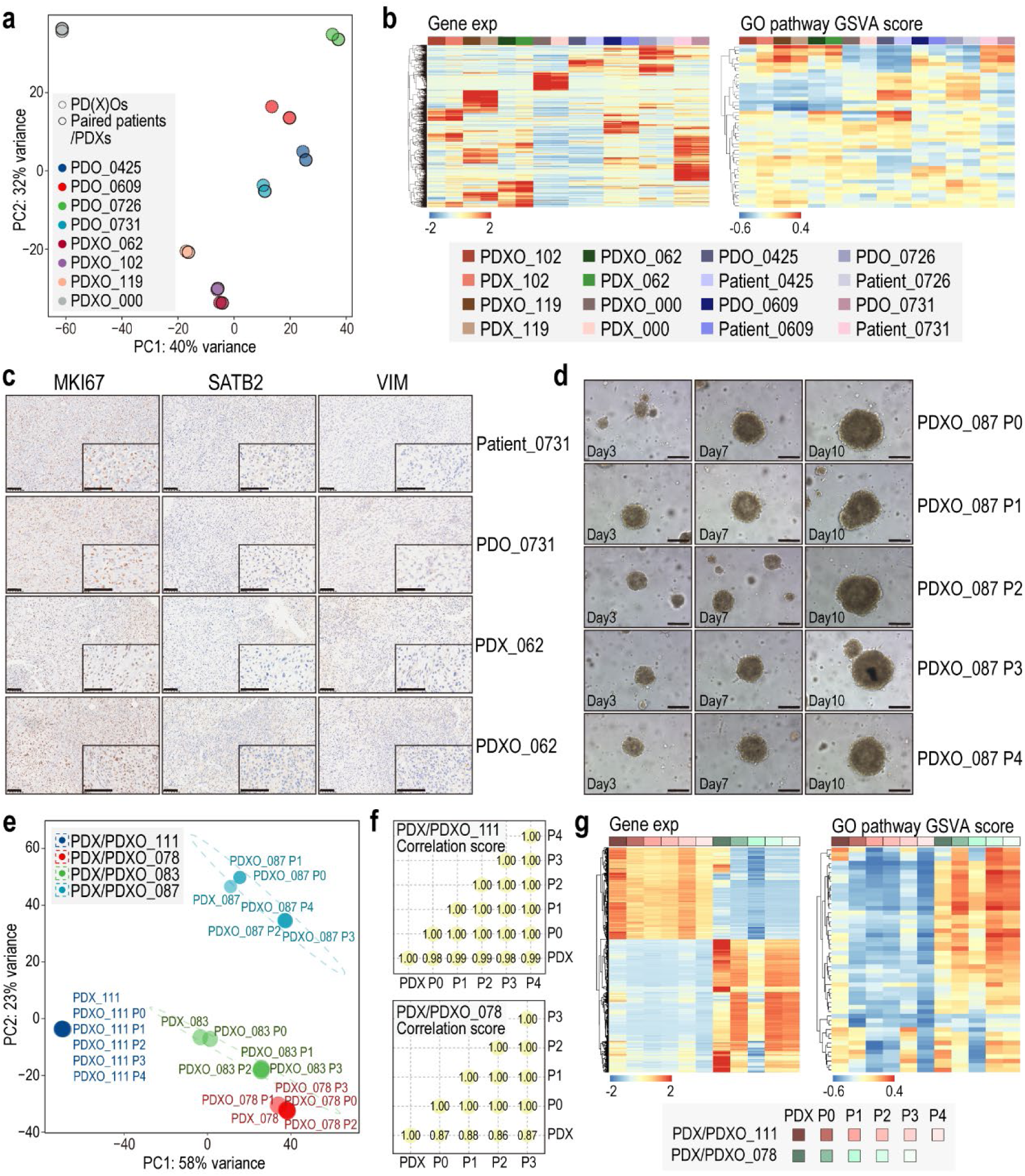
OS PD(X)Os retain inter-heterogeneity of paired patients/PDXs in continuously passages. **(a)** Bulk RNA-seq gene expression principal component analysis (PCA) plots of samples for OS PD(X)O and paired patient/PDX transcriptomes with 90% confidence ellipses. **(b)** Gene expression and gene set variation analysis (GSVA) score heatmap of samples for OS PD(X)O and paired patient/PDX transcriptomes. **(c)** Sample immunohistochemical staining images of MKI67, SATB2 and VIM in OS PD(X)O and paired patient/PDX. Scale bars, 100 μm. **(d)** Sample brightfield images of continuously passaged OS PD(X)Os. Age of OS PDOs in days is listed. Scale bars, 250 μm. **(e)** Bulk RNA-seq gene expression principal component analysis (PCA) plots of samples for continuously passaged OS PD(X)O and paired patient/PDX transcriptomes with 90% confidence ellipses. **(f)** Correlation analysis using Spearman correlation of continuous passaged OS PD(X)O models and paired patient/PDX transcriptomes. **(g)** Gene expression and gene set variation analysis (GSVA) score heatmap of samples for continuously passaged OS PD(X)O and paired patient/PDX transcriptomes. See also **Supplementary Figure 4**.

Further, we investigated the genomic characteristics of OS PD(X)O models in our biobank. Given that OS is characterized by chromosomal copy number variations (CNVs), we initially examined CNVs across different passages of OS PD(X)O models and their paired specimens. Our models exhibited high consistency in CNVs with their paired specimens, and the primary CNV features of OS PD(X)O models were preserved during passaging (**Figure 4a**). Building on our previous study of osteosarcoma multi-omics subtypes1, we further explored alterations in key genes of osteosarcoma. We identified amplifications in critical genes such as MYC, NF1, NOTHC3, VEGFA, RAD21, BRD4, AURKB1, and deletions in key genes including CDKN2A/B, MTAP, RB1, with these CNV features also preserved across passages (**Figure 4b**). Although point mutations are uncommon in osteosarcoma, we also screened for mutations in our biobank and found intriguing and potentially actionable mutations in genes such as BRCA2, KRAS, and SF3B1 (**Figure 4c**). Some mutations may emerge during model establishment or diminish over passages, possibly due to the evolution occurring during tumor cell progression. Additionally, the sensitivity of whole genome sequencing (WGS) should be considered, particularly due to the smaller sample size in iOS models compared to the original tissues. For instance, while KRAS was detected in PDXO_111 P1, it could be detectable by KRAS^G12C^ mutation-enriched probes at P5, with a weakened signal in WGS (**Figure 4d**). Nevertheless, our genomic profiling of OS PD(X)O models demonstrates that our biobank successfully inherits the genetic characteristics of paired tissues from patients/PDXs, reflecting the genomic diversity in the biobank.

**Figure 4.**
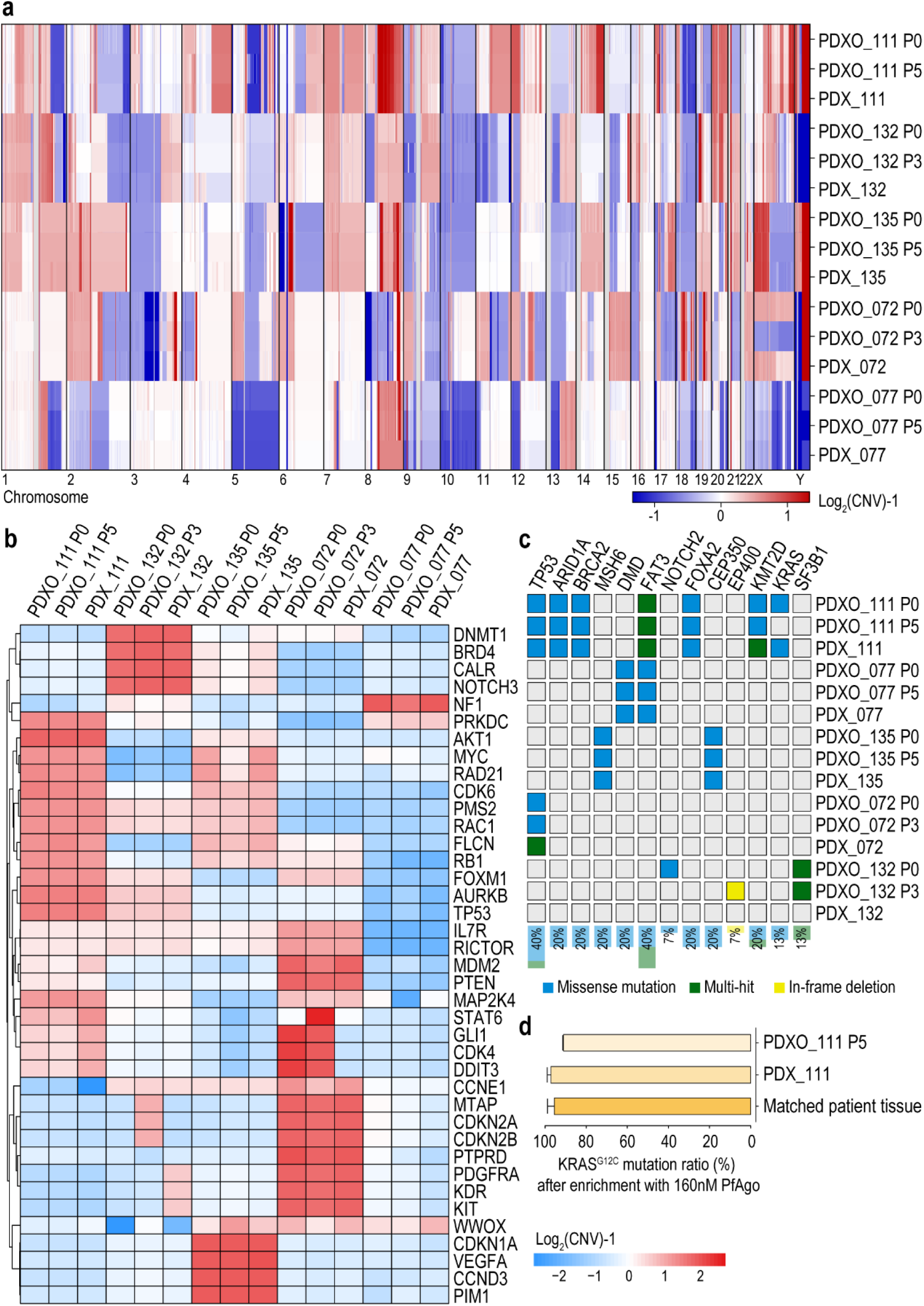
OS PD(X)Os retain inter- and intra-genomic heterogeneity of paired patients/PDXs in continuously passages. **(a and b)** Copy number variations in autosomal chromosomal arms and in osteosarcoma-associated genes identified by whole-genomic sequencing (WGS) of continuous passaged OS PDXO models and paired PDX in our biobank. **(c)** Somatic variants in osteosarcoma-associated genes identified by whole-genomic sequencing (WGS) of continuous passaged OS PDXO models and pairedPDX in our biobank. **(d)** KRAS^G12C^ gene probe assay using 160nM PfAgo. KRAS^G12C^ abundance in PDXO_111 and paired PDX/patient tissue. Values represent mean ± SD (n = 3).

### Establishment of an immune-featured osteosarcoma patient/PDX derived organoid (iOS) model

After confirming that the OS PD(X)O biobank retains the molecular and genomic diversity of OS patients/PDXs, our biobank can serve as a foundation for personalized therapy in OS. However, it lacks elements that simulate patient lymphocytes, especially T lymphocytes that reflect the efficacy of immune checkpoint inhibitors. Therefore, we constructed an **immune-featured osteosarcoma patient/PDX derived organoid (iOS)** model, considering the young age of OS patients and other clinical factors. To ensure procedural consistency, we opted for commercially available, mature human PBMCs as substitutes for patient-derived PBMCs (**Figure 5a**). We endowed the iOS models in our biobank with lymphocyte characteristics, theoretically enabling the iOS to exhibit tumor immune-killing effects. The iOS models can still form spheroids, but there will be some free lymphocytes surrounding them (**Figure 5b**). Using flow cytometry, we detected the distribution of lymphocytes in iOS models, where CD8^+^ T lymphocytes constitute approximately 10% of CD45^+^CD3^+^ T lymphocytes. CD4^+^ T lymphocytes outnumber CD8^+^ T lymphocytes, but approximately half of the CD45^+^CD3^+^ lymphocytes remain undifferentiated, indicating that these cells are still in an immature state without differentiation (**Figure 5c and 5d**). These results provide initial evidence of the successful establishment of iOS models.

**Figure 5.**
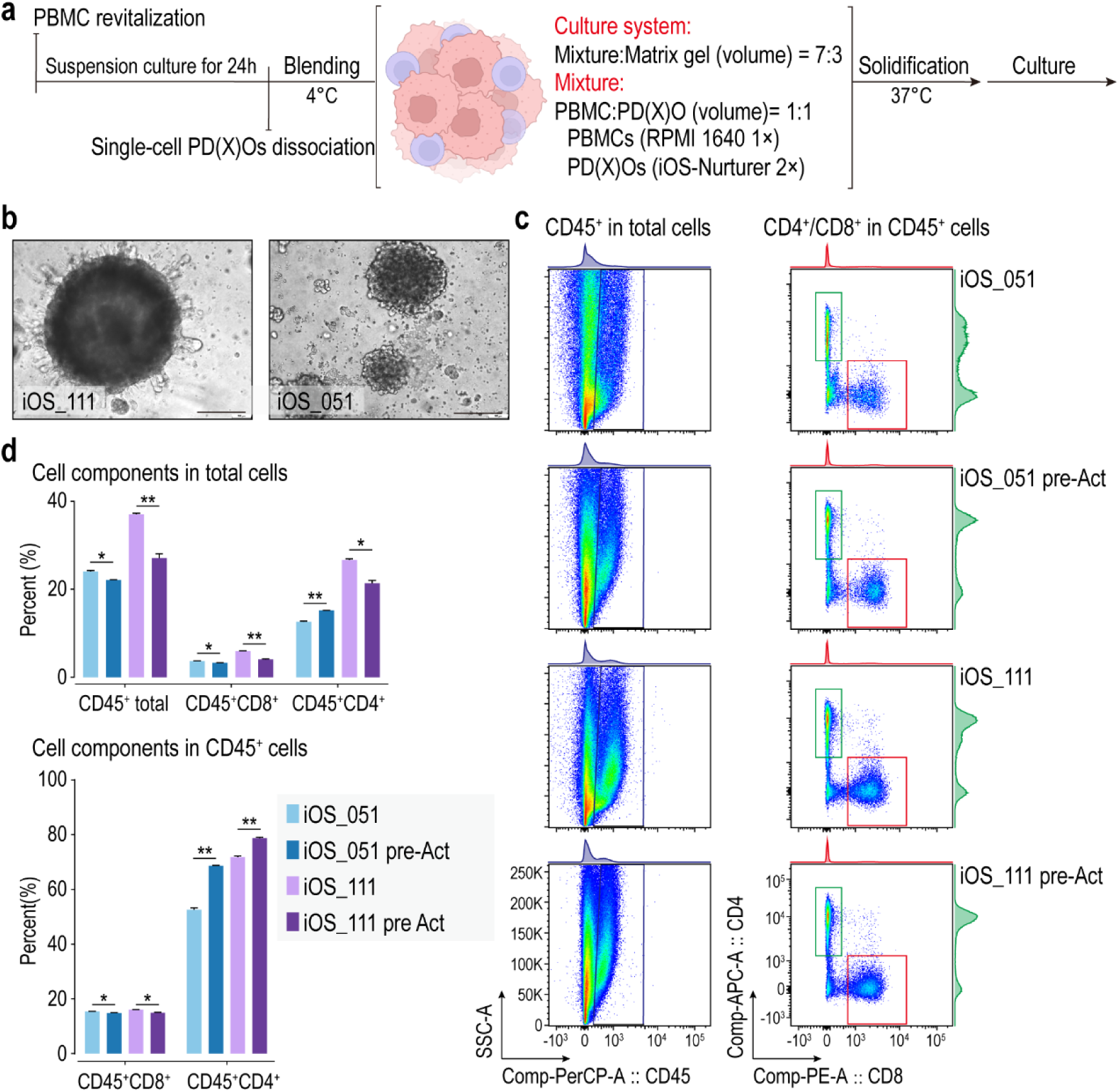
Immune-featured osteosarcoma patient/PDX derived organoid (iOS) models regain T lymphocyte features. **(a and b)** A schematic of the procedure for iOS model establishment with sample brightfield images. Scale bars, 250 μm. (c) Flow cytometry analysis showing the exhibition of CD8^+^ or CD4^+^ T lymphocytes in iOS models, either using pre-activated (pre-Act) or non-activated PBMCs. (d) Quantification of CD45^+^, CD45^+^CD8^+^ and CD45^+^CD4^+^ T lymphocytes in total cells and quantification of CD45^+^CD8^+^ and CD45^+^CD4^+^ T lymphocytes in CD45^+^ total lymphocytes, either using pre-activated (pre-Act) or non-activated PBMCs. Values represent mean ± SD (n = 3).

To validate the biological functionality of iOS models and their consistency with clinical patient treatments, we first assessed the chemosensitivity of iOS models. To validate the biological functionality of iOS models and their consistency with clinical patient treatments, we first assessed the chemotherapy sensitivity of the iOS models. We treated iOS models and their paired OS PD(X)O models with the first-line chemotherapy drug for osteosarcoma, doxorubicin (DOX), and compared the results with the standard chemotherapy effects observed in corresponding patients. Patient_072 did not exhibit lung metastasis during standard chemotherapy (**Figure 6a**), and both iOS_072 and PDXO_072 showed sensitivity to DOX (**Figure 6b**), with noticeable shrinkage at 0.1μM DOX and significant shrinkage at 10μM DOX (**Figure 6c**). For Patient_051, he had already developed lung metastasis when standard chemotherapy began, but there was no increase observed in the early follow-up period (**Figure 6d**). His iOS_051 and PDXO_051 showed sensitivity to DOX (**Figure 6e and 6f**). But for Patient_111, during standard chemotherapy, he developed lung metastases unfortunately (**Figure 6g**). His paired iOS_111 and PDXO_111 models were also insensitive to chemotherapy drugs (**Figure 6h and 6i**). These results indicate that our iOS models constructed from OS PD(X)Os in our OS PD(X)O biobank correlate well with actual clinical treatments for patients. Both iOS and OS PD(X)O models show no difference in reflecting sensitivity to standard chemotherapy.

**Figure 6.**
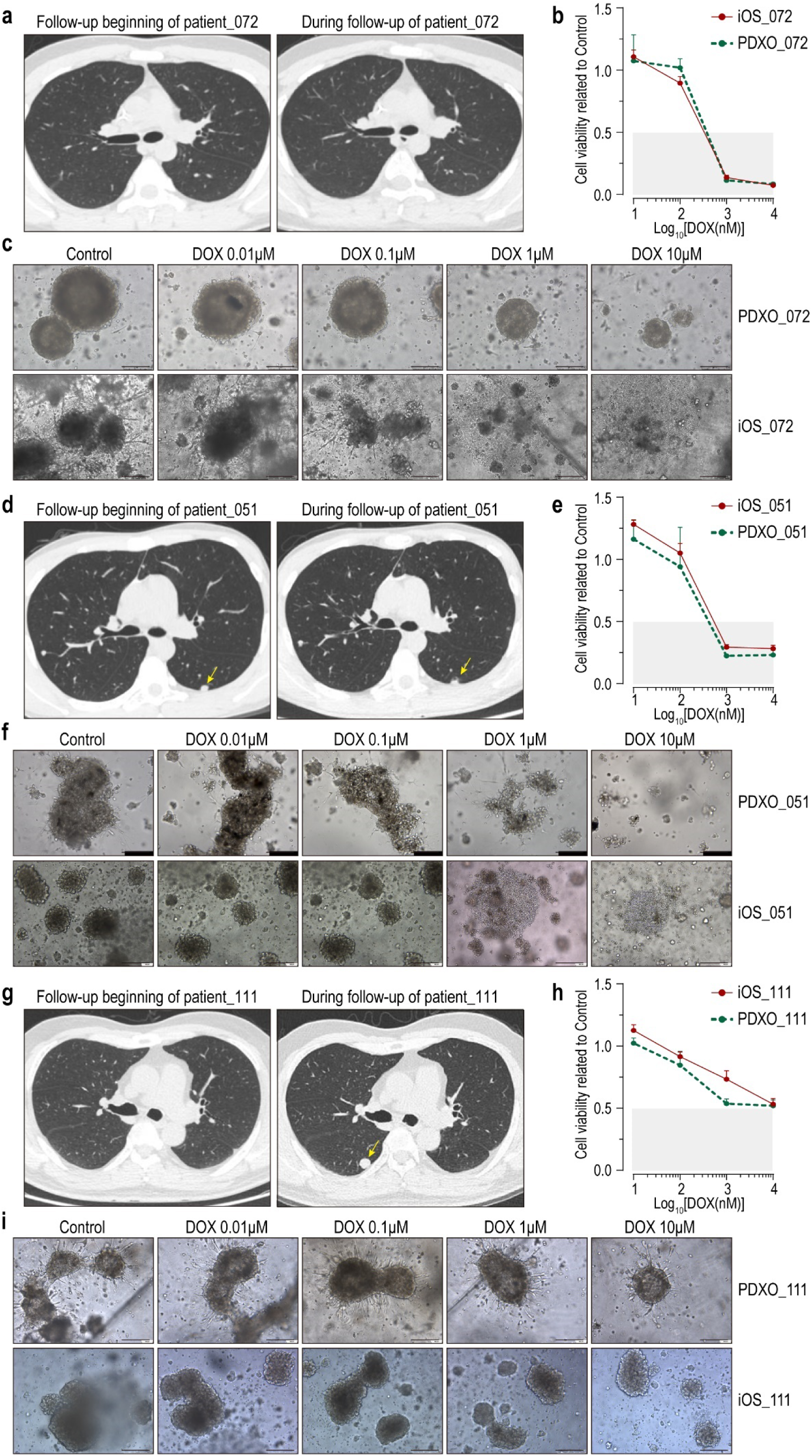
Chemotherapeutic responses of iOS models *in vitro* and retrospective following-up of chemotherapeutic responses of paired patients. **(a, d and g)** Radiological images of patients at the beginning of and during the post-surgery chemotherapy by retrospective following-up. Arrows indicate the OS metastasis lesions. **(b, e and h)** 3D cell viability assay. Cell viability quantifications of iOS and its paired OS PD(X)O models, corresponding for different culture days, treated with Doxorubicin (DOX). Values represent mean ± SD (n > 3). **(c, f and i)** Sample brightfield images of iOS and its paired OS PD(X)O models, treated with Doxorubicin (DOX). Age of OS PDOs in days is listed. Scale bars, 250 μm.

We further assessed the immunobiological functions of the iOS model, particularly the cytotoxic T lymphocytes (CD8^+^ T lymphocytes) functions. Starting from the second day after constructing the iOS model, we administered PD-1 monoclonal antibody every two days, coinciding with the media change schedule of the iOS model (**Figure 7a**). We observed the biological functions and cell activities of the iOS model (**Figure 7b**). We selected iOS_111 and paired PDXO_111 as a case due to its poor response to chemotherapy. Compared to PDXO_111, iOS_111 showed a certain reduction in tumor size, which was statistically significant but did not significantly affect tumor killing, even after PD-1 monoclonal antibody treatment (**Figure 7c**). Considering that iOS model theoretically mimics the real state of patients, our results can be explained, as the overall efficacy of PD-1 monoclonal antibody treatment alone or combined in clinical settings is not ideal ^28–30^. We verified PD-1 expression on CD8^+^ T lymphocytes using flow cytometry, showing a decrease in PD-1 expression on CD8^+^ T lymphocytes after PD-1 monoclonal antibody treatment (**Figure 7d and 7f**). These results indicate the effectiveness of PD-1 monoclonal antibodies on the iOS model, but further personalized treatment research is needed to enhance the cytotoxic effects of CD8^+^ T lymphocytes.

**Figure 7.**
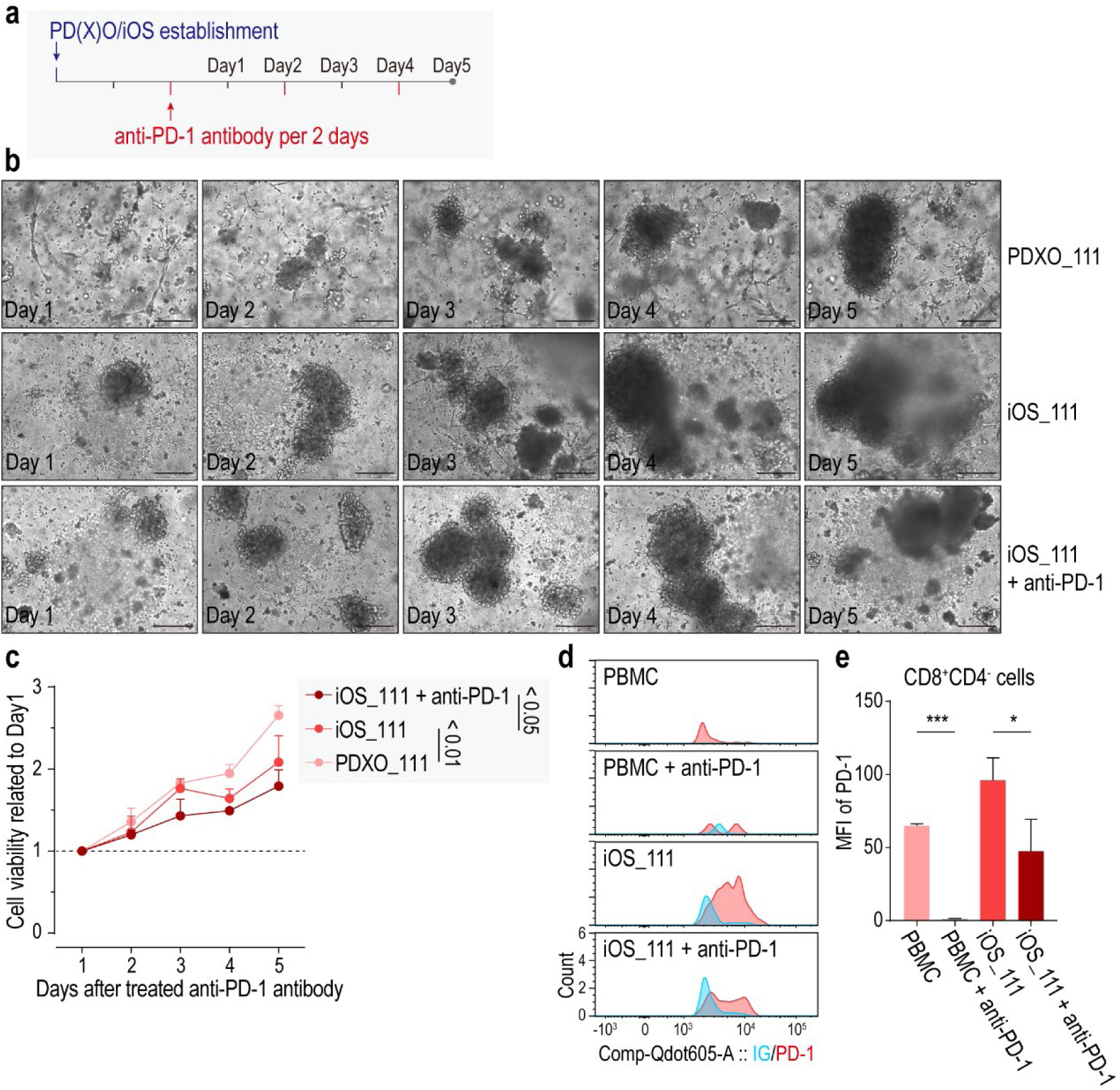
Modeling immunotherapy testing using iOS models treated anti-PD-1 antibody. (a) A schematic of the procedure for iOS model treated with anti-PD-1 antibody. (b) Sample brightfield images of the iOS model, treated with anti-PD-1 antibody (anti-PD-1). Age of OS PDOs in days is listed. Scale bars, 250 μm. (c) 3D cell viability assay. Cell viability quantifications of the iOS model, corresponding for different culture days, treated with anti-PD-1 antibody (anti-PD-1). Values represent mean ± SD (n = 3). (d) Flow cytometry analysis showing the PD-1 expression on CD8^+^ T lymphocytes either in iOS models or PBMCs alone, treated with anti-PD-1 antibody (anti-PD-1). (e) Quantification of the PD-1 expression on CD8^+^ T lymphocytes either in iOS models or PBMCs alone, treated with anti-PD-1 antibody (anti-PD-1). Values represent mean ± SD (n = 3). *** p value < 0.001, * p value < 0.05.

### Personalized drug evaluation utilizing immune-featured osteosarcoma patient/PDX derived organoid models

To validate that the iOS models constructed in this study can guide personalized drug development and application, we used iOS_111 as a specific case and employed a novel PRMT5^MTA^ inhibitor APRN2169 in combination with the immune checkpoint inhibitor to simulate the drug response of patient to their combined application. iOS_111 is derived from PDXO_111, which inherits the genetic characteristics of OS patient Patient_111, namely the loss of genomic locus 9p21.3, including CDKN2A/B and MTAP, and KRAS^G12C^ mutation. Previous studies have reported that tumors harboring MTAP deletion and KRAS mutation exhibit synthetic lethality with MAT2A/PRMT5^31–33^. Inhibition of PRMT5 has been found to enhance tumor sensitivity to immunotherapy^34^, but due to the critical role of PRMT5 protein in T cell development^35,36^, PRMT5 inhibitors exert significant side effects on normal cells or tissues, such as T lymphocytes. Previous results above suggest that combination with APRN2169 may enhance the sensitivity of iOS_111 to immunotherapy. Furthermore, PRMT5^MTA^ inhibitor APRN2169 inhibits activation of MTA-cooperative PRMT5 in only cells carried MTAP deletion but not in normal cells, which makes PRMT5^MTA^ inhibitor APRN2169 more favorable for T lymphocytes within tumor tissue. We first validate the lethality against MTAP deleted osteosarcoma cells, utilizing one constructed MTAP deleted mouse-derived osteosarcoma cell line DuNN MTAP-del in our previous study. The efficacy of APRN2169 in DuNN MTAP-del *in vivo* was confirmed (**Supplementary Figure 5a and 5b**). Combined application of APRN2169 enhanced the sensitivity of DuNN MTAP-del to immune checkpoint inhibitors *in vivo* (**Supplementary Figure 5c and 5d**).

After initially determining the effectiveness of APRN2169, we further validated whether our iOS model can detect the efficacy of APRN2169 combined with immunotherapy. Similar to the use of PD-1 monoclonal antibody alone, APRN2169 or combined with PD-1 monoclonal antibody was administered starting two days after iOS model establishment, with drug administration every two days coinciding with medium changes (**Figure 8a**). After 5 days of drug treatment, consistent results showed statistically significant differences between both mono-therapy groups (anti-PD-1 or APRN2169) and the control group (Control), as well as between the combined therapy group (Combined) and Control. Tumor cell activity was significantly reduced in Combined, as observed qualitatively and quantitatively (**Figure 8b and 8c**). However, APRN2169 combined with PD-1 monoclonal antibody was not did not show efficacy in the CDKN2A/MTAP wildtype iOS model, such as iOS_135 (**Supplementary Figure 6a and 6b**). In addition, to demonstrate the efficacy of combination therapy, flow cytometry was used to assess the distribution and content of CD8^+^ T lymphocytes. In Combined, there was a statistically significant increase in CD8^+^ T lymphocytes (**Figure 8d and 8e**). Unexpectedly, the group treated with APRN2169 alone exhibited the lowest PD-1 expression on CD8^+^ T lymphocytes. Whatever, both the single-agent and combination therapy groups showed significantly reduced PD-1 expression on CD8^+^ T lymphocytes compared to Control (**Figure 8f**). The above results suggest that our iOS model can serve as an effective model for personalized drug development in osteosarcoma treatment.

**Figure 8.**
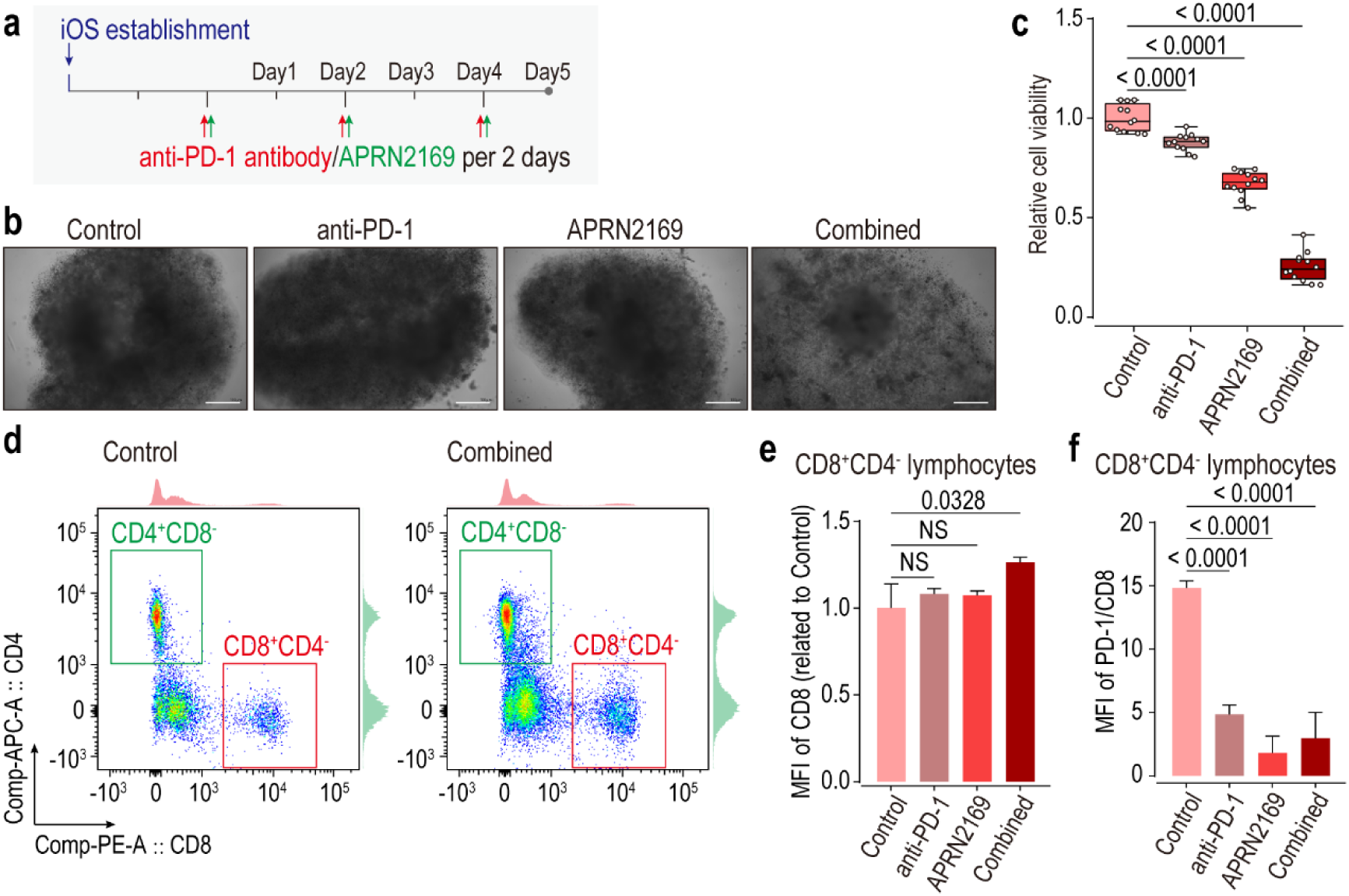
Personalized immunotherapy testing using MTAP-deleted iOS model treated with PRMT^MTA^ inhibitor APRN2169 and anti-PD-1 antibody combination. **(a)** A schematic of the procedure for iOS model treated with anti-PD-1 antibody and PRMT5^MTA^ inhibitor APRN2169 combination. **(b)** Sample brightfield images of the iOS model, treated with anti-PD-1 antibody and PRMT5^MTA^ inhibitor APRN2169 combination. Age of iOS model is at Day5 after treated. Scale bars, 100 μm. **(c)** 3D cell viability assay. Cell viability quantifications of the iOS model, treated with anti-PD-1 antibody and PRMT5^MTA^ inhibitor APRN2169 combination. Age of iOS model is at Day5 after treated. Values represent mean ± SD (n = 12). **(d)** Flow cytometry analysis showing the exhibition of CD8^+^ or CD4^+^ T lymphocytes in iOS models, treated with anti-PD-1 antibody and PRMT5^MTA^ inhibitor APRN2169 combination. (**e and f**) Quantification of the CD8 or PD-1 expression on T lymphocytes in iOS models, treated with anti-PD-1 antibody and PRMT5^MTA^ inhibitor APRN2169 combination. Values represent mean ± SD (n = 3). See also **Supplementary Figure 5** and **Supplementary Figure 6.**

## Discussion

Our OS PD(X)O models and biobank recapitulate the paired patient/PDX heterogeneity as evidenced by (1) histological staining illustrating similar tissue architecture and cellular morphologies, (2) immune-histological staining showing a similar performance on common biomarkers, (3) bulk RNA transcriptomes identifying maintenance of biological function signatures and (4) genomics confirming the inheritance of CNVs and somatic variants. Our xenografts of OS PD(X)O models displayed efficient engraftment. Additionally, our iOS models regain the T lymphocyte populations for recapitulate a complete tumor microenvironment of OS. We demonstrate that the iOS models could simulate the chemotherapy responses of paired patients by retrospective following-up. We also demonstrate iOS models show response to immune checkpoint therapy. Based on typical specific genomic mutations, iOS models show differential responses to targeted drug mono-treatment and combined with immunotherapy. By employing iOS models carried MTAP deletion and KRAS mutation, we identify the efficacy of a novel PRMT5^MTA^ inhibitor APRN2169 and combination with PD-1 monoclonal antibody. Our method and models give it possible to accelerate personalized clinical or pre-clinical strategies in 7 to 10 days and influence clinical decisions on diagnoses and treatments. Our models and biobank of OS could be as a resource for future biological studies, therapeutic high-throughput drug screenings and even clinical trials.

### OS Models and Biobank Inheriting Patient Heterogeneity Suitable to Clinically Rapid Timing

One important feature of our models and their generation methods are that through single-cell dissociation, impurities like small bone fragments are removed, and the cells autonomously reassemble to regain their cell-cell morphologies and structures. The entire culture cycle lasts 7 to 10 days, demonstrating high intra-group consistency and stability. With the passaging and cryopreservation methods developed in this study, the cultures can be continuously maintained and preserved. Given that osteosarcoma patients often have a short survival period after recurrence or metastasis, the timeframe for monitoring personalized therapeutic drugs is critical. Meanwhile, for rare diseases characterized by complex genetic backgrounds and significant individual variations, we contend that relying on large-scale databases to identify common drugs for clinical translation carries inherent risks. It is imperative to swiftly customize diagnostics and treatments for each patient to accurately pinpoint effective therapeutic agents. Even patient-derived xenograft (PDX) models^37,38^, which can replicate individualized genetic backgrounds, may struggle to keep pace in rapid treatment cycles. Thus, the development of PD(X)O models is crucial for diseases like osteosarcoma. As the tumor collection in SGH-OS cohort and model generation is undergoing, we believe that our OS PD(X)O models and biobank will be a useful resource for future biological studies, therapeutic high-throughput drug screenings and even clinical trials.

### Establishment of Osteosarcoma Models Available for Research of Tumor Immunology

Our study established an iOS model not only to screen targeted drugs but also to facilitate various immunotherapies, including immune checkpoint inhibitors, cell therapies, and oncolytic viruses. Our iOS model re-gain the T lymphocyte features of OS microenvironment and could be used for development of personalized immune therapeutic strategies. For recurrence and metastasis OS patients, they typically face a challenging prognosis, and unfortunately, the available treatment options to enhance their survival are limited. Given the successes in immunotherapy for challenging solid tumors, but immunotherapy, especially immune checkpoint therapy (ICT), alone or in combination, failed to achieve the predefined target on OS ^28,30^. Unlike Ewing sarcoma or liposarcoma, OS exhibits a deficiency in tumor-infiltrating lymphocytes (TILs)^29^. Consequently, models unlike traditional *in vitro* models (such as K7/K7M2 or DuNN^39,40^ mouse osteosarcoma cell lines) inheriting patient heterogeneity and recapitulating activation of anti-tumor immune urgently needs to be developed, for elucidating the biological mechanisms underlying the recruitment and activation of tumor-infiltrating lymphocytes (TILs) in the OS tumor microenvironment (TME) and identifying potential intervention targets to enhance the therapeutic efficacy of OS immunotherapy. In previous studies, immunogenomic characteristics, such as CDKN2A/MTAP deletion, may decrease the abundance of tumor-infiltrating lymphocytes (TILs) and contribute to primary resistance to ICT ^41,42^. CDKN2A/MTAP deletion is also a common genomic mutation in OS ^4,43^, and it might be a potential target for enhancing immune therapy responses of OS patients. Fortunately, in our biobank, we have models carried CDKN2A/MTAP deletion and we employed MTAP deletion iOS models and MTAP wildtype models to testing the efficacy of a novel PRMT5^MTA^ inhibitor APRN2169. APRN2169 shows selective efficacy on MTAP deletion iOS models but not on MTAP wildtype iOS models. We believe our iOS models can better support novel drug development and play a significant role in advancing immunotherapy or treatments targeting the tumor microenvironment.

### Practical Limitations and Future Applications

Nonetheless, our findings reveal that the iOS model can only simulate patient responses over a limited period. We hypothesize that in the short term, the iOS model mirrors the evolution of patient tumor tissue. However, ongoing changes necessitate adapting the iOS model longitudinally to align with evolving therapeutic needs. For instance, PDXO_051 initially matched the patient’s treatment response despite inheriting most patient-derived genetic traits. However, approximately six months later, the patient succumbed to disease progression. As we embarked on constructing the iOS model, our process and objective involved disaggregating osteosarcoma tissue into single cells and allowing cells to self-assemble into a matrix using only provided raw materials. During this process, we observed scattered tumor cells around the iOS model forming sporadic satellite foci, suggesting mutual cellular attraction (data not shown). Subsequent verification will utilize continuous bright-field imaging. Within our biobank, we observed diverse biological behaviors of fibroblasts. Interestingly, at times, our models exhibits pulsatile movements akin to a heartbeat (**Supplementary Video 1**). Occasionally, tumor-associated fibroblasts gradually communicate with each satellite focus during the formation of the iOS model, integrating them into a cohesive whole. These behaviors may be linked to the invasiveness of osteosarcoma and warrant further detailed investigation.

Our iOS model still has many areas that need improvement. According to theory, PDX models can be established due to the lack of tumor-killing T cell function in immunodeficient mice. However, macrophages and fibroblasts, for example, participate in the formation of tumor tissue ^44^. This is why we initially focused on restoring lymphocyte characteristics in the iOS model. We have only assessed the functionality of CD8^+^ T lymphocytes, and the comprehensive functionality of other cell components in the iOS model remains a significant undertaking that we plan to refine in future experiments. Additionally, our study did not investigate materials like calcium chloride, disodium glycerophosphate, etc., which serve as osteogenic materials for osteoblasts. We only added collagen, which may suffice in the short term but has not yet achieved our ideal endpoint for osteosarcoma or bone organoid models.

In summary, our immune-featured iOS model recapitulates the patient heterogeneity and typical biological and genomic characteristics of OS. Based on our models, our biobank has the potential for clinical timely drug testing of personalized therapy strategies and clinical decisions, with broad applications in basic, translational even clinical researches of OS.

### Supporting information

Supplementary figures

Supplementary Table 1

Supplementary Video 1

## Declaration

All authors declare no conflict of interest.

## Ethical Approval

Our study was approved by the Institutional Research Ethics Committee of Shanghai General Hospital, Shanghai Jiao Tong University School of Medicine (2021KY103).

## Author information

### Contributions

Y.Q. Hua, L. Yang, W. Sun and Z.D. Cai designed the research. H.R Mu and Y.N. Tao wrote the manuscript. H.R Mu, D.Q. Zuo, H.S. Wang, J.K. Shen, M.X. Sun, Y.Z. Liang, H.Y. Wang and X.Y. Yang collected the clinical data and specimens. H.R. Mu, Y.N. Tao, Q. Zhang, X. He and L.Y. Zhang performed the basic experiments. J.Z Wang, Y.N. Tao, Q. Zhang and H.R. Mu performed bio-informatic analysis. H.R. Mu, Y.N. Tao, X.Y. Yang, X. He, Q. Zhang and X.M. Jin established the culture method for OS PD(X)O, further iOS model and biobank in the research. H.R. Mu, Y.Y. Gong, W.X. Chen and Y.N. Tao established the dissociation method for OS PD(X)O, further iOS model and biobank in the research. B. Yao, W. Mao, F.J. Asward, J.D. Joseph and M.X. Li provided the compound APRN2169 and its methods of administration, including the route of administration, dosage, and formulation method. Y.Q. Yang, M. Jiao and Y.H. Jin established the PDX models in the research. S. Ding and Q. Liu performed the KRAS^G12C^ gene probe assay. T. Zhang, J. Xu and J. Lin supervise the experiments. Z.Y. Wang and L. Lv managed the reagents and expenses used in the project. All authors have read and approved the article.

### Corresponding author

Correspondence to Zhengdong Cai, Liu Yang, Wei Sun and Yingqi Hua.

### Availability of data and materials

The data that support the findings of this study are available on request from the corresponding authors upon reasonable request.

## Acknowledgements

We would be deeply grateful to Professor Bing Li [*Shanghai Jiao Tong University School of Medicine*, China] for his help that greatly improved this work. We would be deeply grateful to Jun Zhang [*SeekGene*, China] and Chen Chen [*Benagen*, China] for their suggestions and greatly help to improve this work. We would be deeply grateful to Professor Tianlong Li [Harbin Institute of Technology, China], Zhanxiang Zhang [Harbin Institute of Technology, China] and Chenlu Liu [Harbin Institute of Technology, China] for the help of 3D printing at the beginning of this research though we finally did not use the 3D printing architecture.

