## Supplementary figures for "A Self-Assembling Immune-Featured Osteosarcoma Patient/PDX Derived Organoid Model and Biobank for Personalized Immune Therapy"

a

| Example | Cell viability (%) | Total cells | Live cells | Died cells | Average diameter (μm) | Average roundness | Aggregation rate (%) |
|---|---|---|---|---|---|---|---|
| | | Conc./mL | Conc./mL | Conc./mL | | | |
| Example_1 | 100 | 356000.00 | 356000.00 | 0.00 | 16.33 | 0.9 | 7.4 |
| Example_2 | 98.46 | 880000.00 | 866000.00 | 13500.00 | 14.45 | 0.87 | 8.98 |
| Example_3 | 97.44 | 995000.00 | 970000.00 | 25500.00 | 14.78 | 0.82 | 12.56 |
| Example_4 | 97.26 | 837000.00 | 815000.00 | 22900.00 | 14.07 | 0.81 | 15.56 |
| Example_5 | 98 | 1220000.00 | 1190000.00 | 24400.00 | 14.63 | 0.74 | 24.53 |
| Example_6 | 95.15 | 946000.00 | 901000.00 | 45900.00 | 12.94 | 0.84 | 14.47 |
| Example_7 | 99.39 | 944000.00 | 938000.00 | 5740.00 | 12.34 | 0.92 | 6.32 |
| Example_8 | 98.26 | 1120000.00 | 1100000.00 | 19500.00 | 11.27 | 0.91 | 6.23 |

b

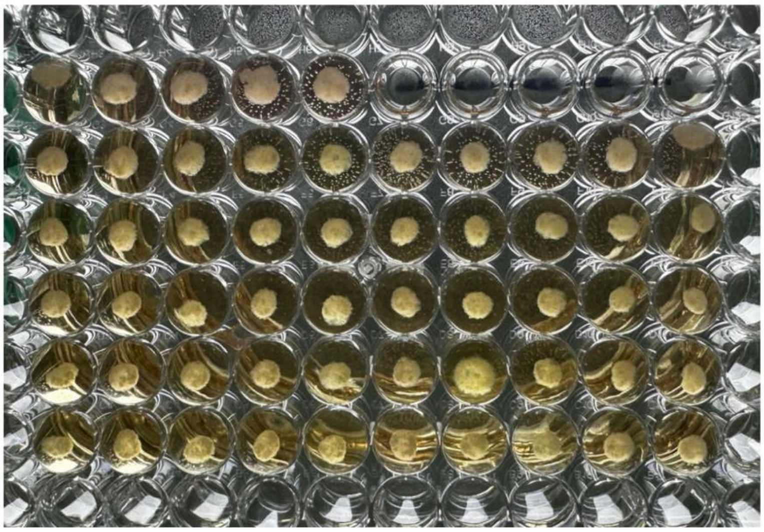

**Supplementary Figure 1** Supplement of figure 1.

(**a**) Information of single-cell OS PD(X)O model suspensions. (**b**) Appearance of OS PD(X)O models in 96-well plates.

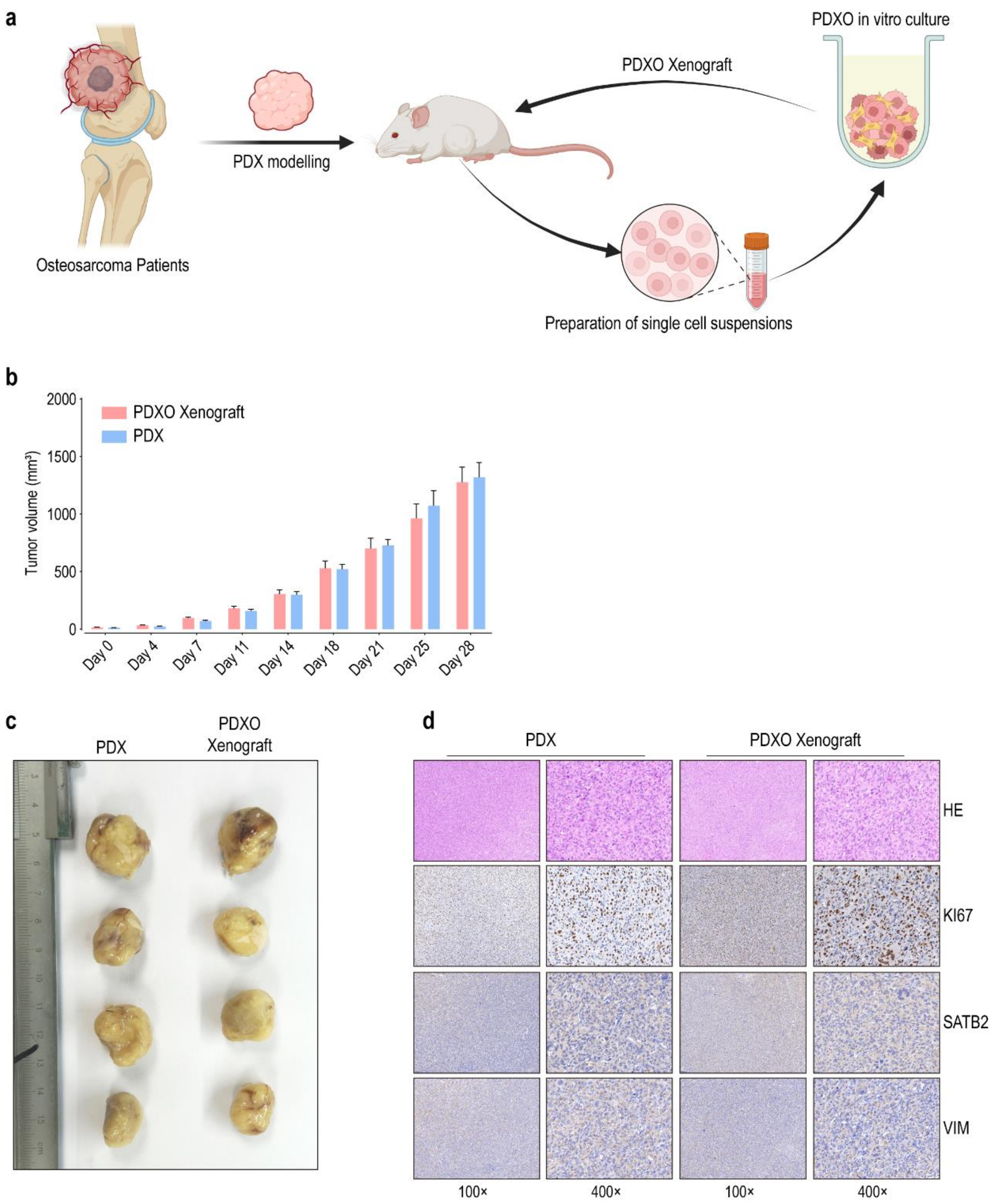


**Supplementary Figure 2** Supplement of figure 1.

(**a**) Flowchart of PD(X)O xenograft establishment. (**b** and **c**) Growth and gross specimens of PD(X)O xenograft and paired PDX, PDXO_111 for example. (**d**) Pathological and typical biological features of PD(X)O xenograft and paired PDX.

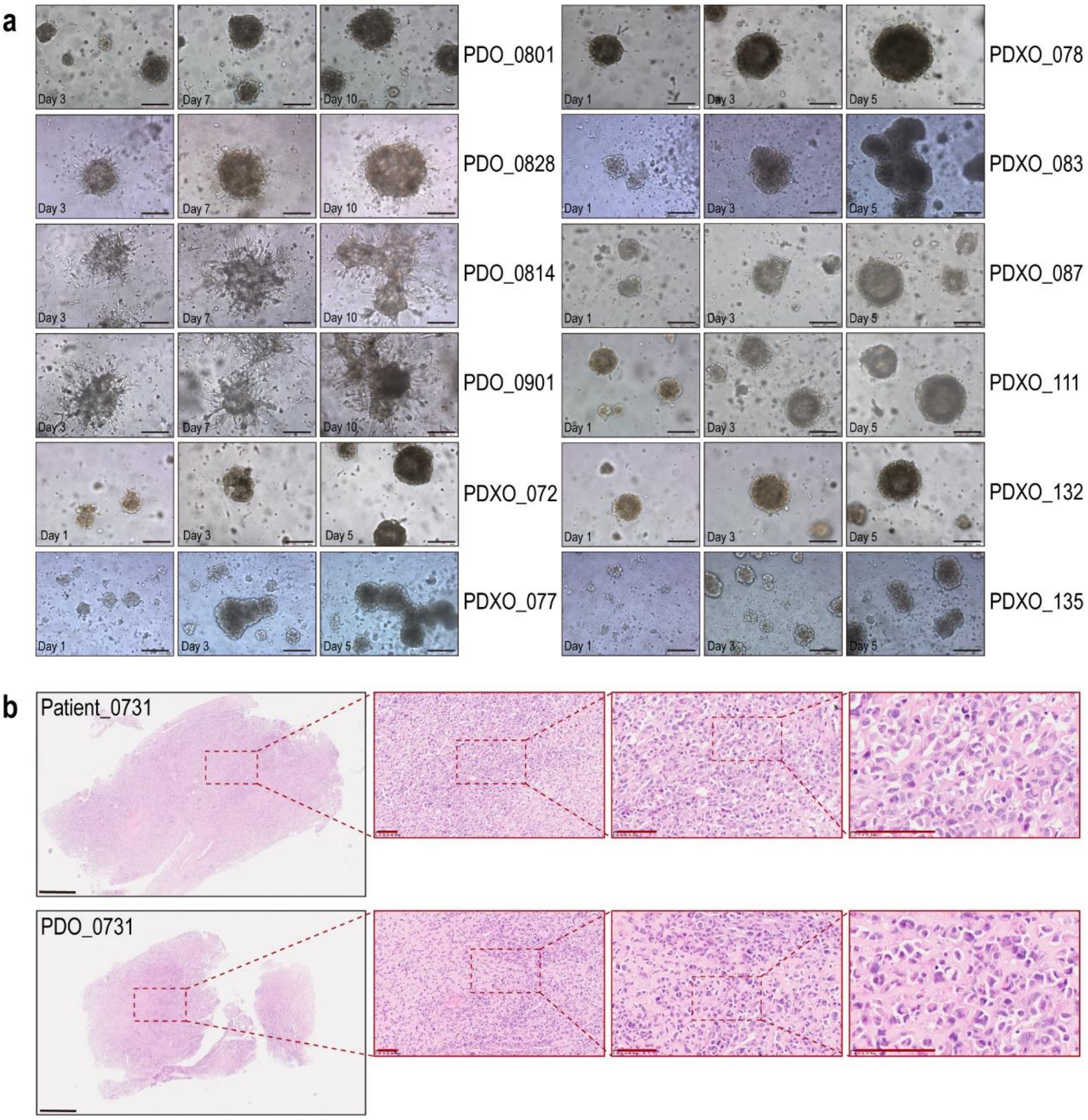


**Supplementary Figure 3** Supplement of figure 2.

(**a**) Additional data of figure 2b and 2d. (**b**) Additional data of figure 2f.

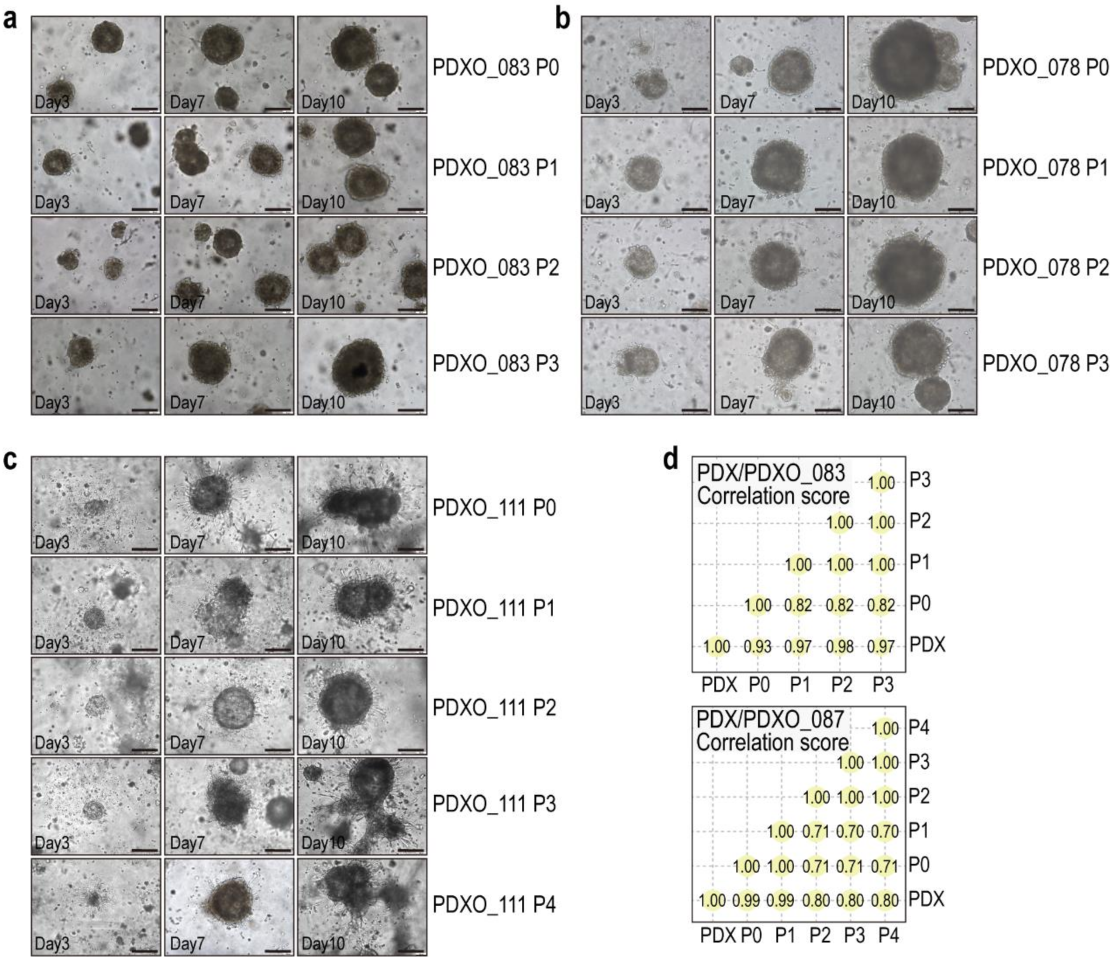


**Supplementary Figure 4** Supplement of figure 3.

(**a**, **b** and **c**) Additional data of figure 3d. (**d**) Additional data of figure 3f.

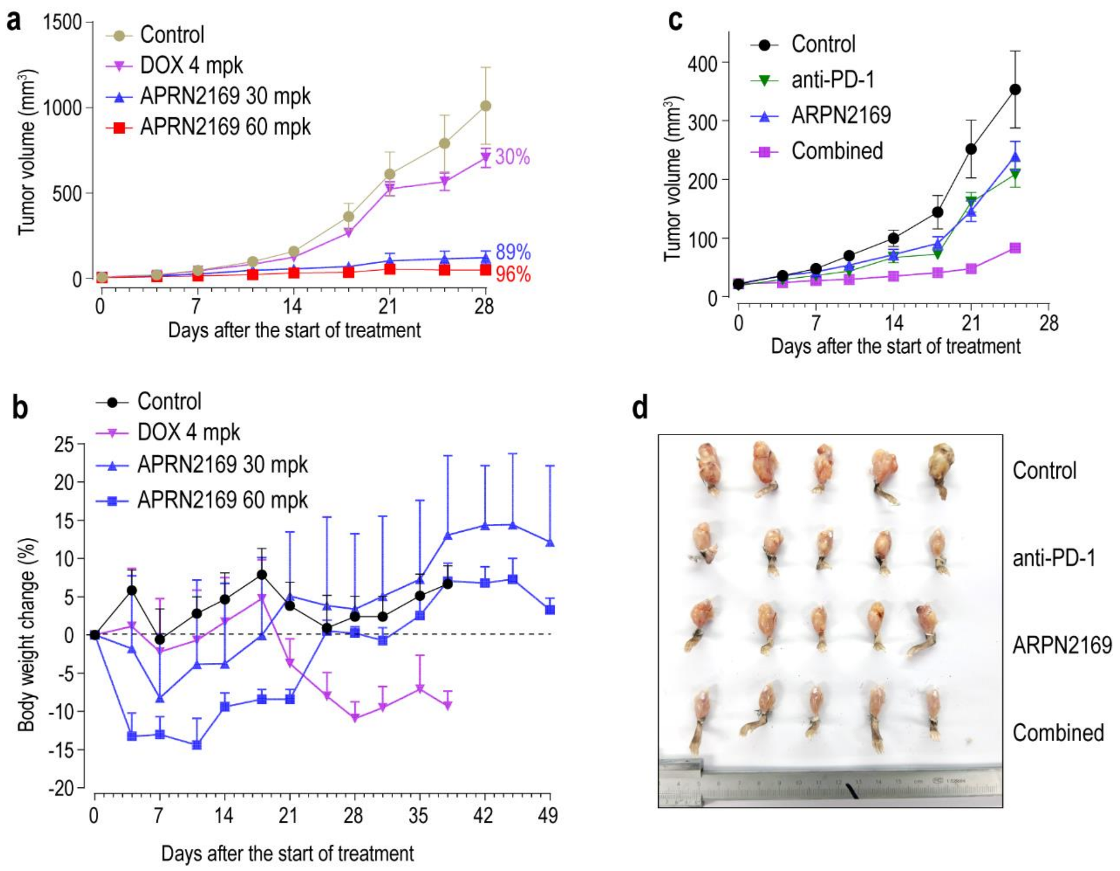


**Supplementary Figure 5** Supplement of figure 8.

(**a** and **b**) Growth and body weight change of DuNN MTAP-del mouse xenografts treated APRN2169 compared with treated Doxorubicin (DOX). (**c** and **d**) Growth and gross specimens of DuNN MTAP-del mouse xenografts treated with APRN2169 and anti-PD-1 antibody. mpk, mg/kg.

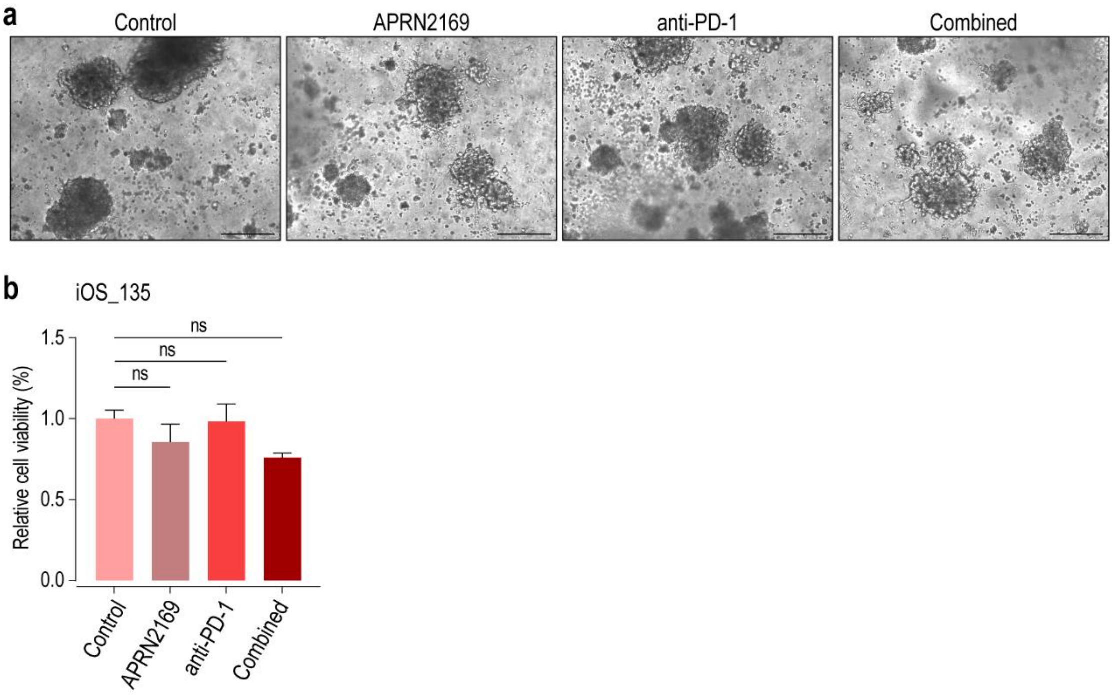


**Supplementary Figure 6** Supplement of figure 8.

**(a)** Sample brightfield images of the iOS model without MTAP deletion, treated with anti-PD-1 antibody and PRMT5$^{MTA}$ inhibitor APRN2169 combination. Age of iOS model is at Day5 after treated. Scale bars, 250 µm. **(b)** 3D cell viability assay. Cell viability quantifications of the iOS model without MTAP deletion, treated with anti-PD-1 antibody and PRMT5$^{MTA}$ inhibitor APRN2169 combination. Age of iOS model is at Day5 after treated. Values represent mean ± SD (n = 3).
